# Investigating the Complexity of Gene Co-expression Estimation for Single-cell Data

**DOI:** 10.1101/2023.01.24.525447

**Authors:** Jiaqi Zhang, Ritambhara Singh

## Abstract

With the rapid advance of single-cell RNA sequencing (scRNA-seq) technology, understanding biological processes at a more refined single-cell level is becoming possible. Gene co-expression estimation is an essential step in this direction. It can annotate functionalities of unknown genes or construct the basis of gene regulatory network inference. This study thoroughly tests the existing gene co-expression estimation methods on simulation datasets with known ground truth co-expression networks. We generate these novel datasets using two simulation processes that use the parameters learned from the experimental data. We demonstrate that these simulations better capture the underlying properties of the real-world single-cell datasets than previously tested simulations for the task. Our performance results on tens of simulated and eight experimental datasets show that all methods produce estimations with a high false discovery rate potentially caused by high-sparsity levels in the data. Finally, we find that commonly used pre-processing approaches, such as normalization and imputation, do not improve the co-expression estimation. Overall, our benchmark setup contributes to the co-expression estimator development, and our study provides valuable insights for the community of single-cell data analyses.

## 1 Introduction

Analyzing the co-variations of gene expression across samples, also known as gene co-expression analysis, is essential for understanding complicated biological processes. For example, gene co-expression analysis can prioritize candidate disease genes (Presson et al., 2008; Van Dam et al., 2018) and molecules of disparate cellular states (Carter et al., 2004), and annotate functionalities of unknown genes (Serin et al., 2016). In previous studies on microarray or bulk RNA-seq data, gene co-expression estimation has shown superior performance for genetic module discovery (Stuart et al., 2003), gene-disease prediction (Van Dam et al., 2018; Langfelder and Horvath, 2008), and biological-processes-related genes detection (Usadel et al., 2009; Aoki et al., 2007). Recently, advances in the high throughput single-cell RNA sequencing (scRNA-seq) technology have enabled researchers to profile gene expression at a single-cell resolution (Hwang et al., 2018; Chen et al., 2019). scRNA-seq datasets allow us to investigate the gene expression of individual cells, revealing heterogeneity among cell populations. These datasets are large, generally consisting of expression for tens of thousands of cells and genes (Macosko et al., 2015; Picelli et al., 2014), thus allowing population-based gene co-expression estimation. However, these new single-cell data opportunities also present new challenges for statistical analysis (Lähnemann et al., 2020). For example, it is notoriously difficult to deal with high sparsity and heterogeneity in scRNA-seq data. Therefore, efficient and robust gene co-expression estimation models for scRNA-seq data are urgently needed in the field.

We formulate the gene co-expression estimation on scRNA-seq data as an unsupervised undirected graph structure learning task (Fig. 1). Concretely, given a tissue/organ with heterogeneous cells, scRNA-seq technique first separates it into individual cells and sequences gene expression at a single-cell level (Haque et al., 2017). The scRNA-seq data can be formalized into an expression matrix 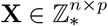 of *n* cells and *p* genes ^2^, where each element **X**_*ij*_ is a count value denoting the expression level of gene *j* at cell *i*. Given the expression matrix, we say a pair of genes are co-expressed if the genes have similar expression patterns. So the goal of gene co-expression estimation is to estimate a symmetric gene similarity matrix **W** ∈ ℝ*p*×*p* with **W**_*jk*_ indicating the similarity score between genes *j* and *k*. We can convert the similarity matrix to an adjacency matrix **G** ∈ {0, 1} ^*p*×*p*^, encoding a co-expression network with nodes denoting genes and undirected edges connecting two nodes *i* and *j* if the corresponding pair of genes are significantly co-expressed, i.e., the similarity score **W**_*ij*_ higher than a user-defined threshold. Estimating the undirected co-expression network is also a fundamental step in directed gene regulatory network (GRN) inference (Nguyen et al., 2021), in which methods can perform enrichment to determine transcript factors and target genes on the undirected network and assign directions to edges. This is suggested by the fact that co-expressed genes are likely to be involved in similar biological functions and share transcription factor binding sites (Marco et al., 2009; Kang et al., 2011; Oldham et al., 2008; Allocco et al., 2004; Yeung et al., 2004; Yin et al., 2021).

**Figure 1:**
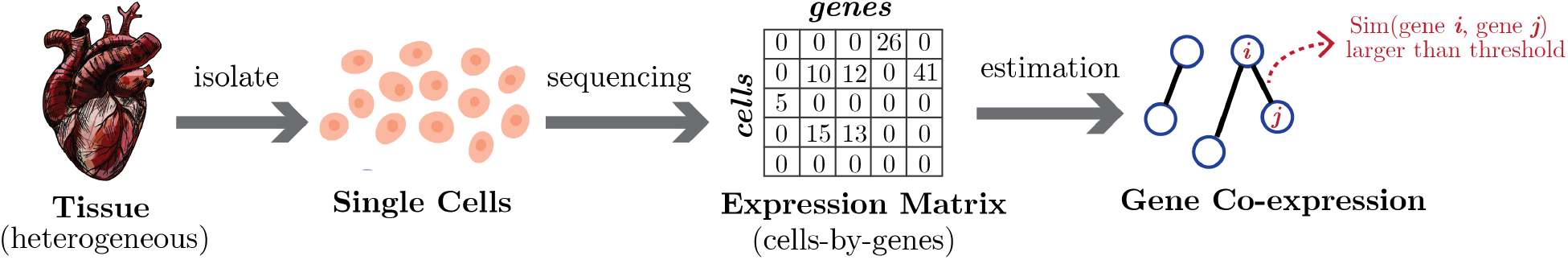
Illustration of gene co-expression estimation on scRNA-seq data. scRNA-seq technique records gene expression of heterogenous cells that are isolated from tissues. The scRNA-seq data is represented as a cell-by-gene matrix, with each matrix element a count value denoting the gene’s expression level at a cell. The expression matrix is then used to estimate gene co-expression, which encodes a graph structure with nodes indicating genes and edges denoting co-expression.

Previous methods have proposed various measurements for computing the gene similarity score and constructing the co-expression network. They generally fall into three categories: (1) The first group evaluates the co-expression between genes by the correlation coefficients of gene expressions. For example, (Langfelder and Horvath, 2008) proposed Weighted Correlation Network Analysis (WGCNA) to construct gene co-expression networks through weighted correlation networks. (2) The second group utilizes Mutual Information (MI) as the co-expression measurement. For example, (Butte and Kohane, 1999) estimated MI via histogram or binning technique. (Margolin et al., 2006) computed MI with differential entropy that averaged the log probability of gene marginal distributions. (Faith et al., 2007) estimated the likelihood of MI instead of calculating significance scores as in the previous works. (Meyer et al., 2007) decomposed the network inference into a series of MI-based supervised gene selection procedures. (3) Methods in the last group are based on structural models. For example, (Yin and Li, 2011; Danaher et al., 2014; Taeb et al., 2020) assumed the raw counts or log-transformed gene expression data followed Gaussian(-like) distributions and adopted Gaussian Graphical Model (GMM) (Friedman et al., 2008) to estimate the partial correlation between genes. GENIE3 (Huynh-Thu et al., 2010) decomposed the task into several regression subproblems and solved them with tree-based models. Also, (Allen and Liu, 2013) estimated gene co-expression with local Poisson graphical models.

However, these methods were mainly proposed for microarray and bulk RNA-seq data. The excess zero counts and heterogeneity in scRNA-seq data present several challenges for the gene co-expression estimation task. Some recent works (Iacono et al., 2019; Cha and Lee, 2020; Wu et al., 2021) have tried to directly apply correlation-based methods on scRNA-seq data. But the correlation coefficient is not a robust measurement for very-sparse scRNA-seq data (Sanchez-Taltavull et al., 2020). Moreover, the Gaussian or Poisson data distribution assumptions in these works may be incorrect, because recent studies (Risso et al., 2018; Townes et al., 2019; Svensson, 2020) have shown that scRNA-seq data follow negative binomial or zero-inflated distributions. To enable co-expression estimation for single-cell data, (Chan et al., 2017) proposed Partial Information Decomposition and Context (PIDC), which used partial information decomposition to explore statistical dependencies between genes in scRNA-seq data. scLink (Li and Li, 2021) revised GGM learning by explicitly modeling zeros in scRNA-seq data. Similarly, (Park et al., 2021; Choi et al., 2017) modeled excess zeros in data with (zero-inflated) negative binomial distributions. But in these works, the simulated datasets used for method evaluations are less sparse than the real-world single-cell experimental data. Two simulated datasets in (Li and Li, 2021) and (Choi et al., 2017), have around 50% and 20% zero counts, respectively, but the experimental scRNA-seq data can have more than 80% of zeros (Hicks et al., 2018; Qiu, 2020). Although these methods showed their considerable ability to recover gene co-expression networks on simulated data, the simulated datasets used in these studies did not exhibit the high sparsity level and overdispersed distribution of experimental scRNA-seq data. Evaluations on such simulations conceal the complexity of estimating co-expression networks from sparse single-cell data. Therefore, we need simulations that accurately capture real-world experimental datasets’ underlying properties to build robust methods for the co-expression estimation task.

In this paper, we test state-of-the-art gene co-expression estimation methods on two novel simulation processes, revealing the complexity of co-expression estimation for single-cell data and suggesting potential directions for improvement. First, we apply two novel simulation processes utilizing information learned from real-world single-cell data (Sec. 3.2) to produce more realistic benchmarking schemes. These schemes will assist in more robust method developments for the gene co-expression estimation task. Next, we perform a comprehensive analysis of the existing co-expression estimation methods. We benchmark nine state-of-the-art and representative gene co-expression estimation methods on ten simulated datasets with different settings and eight experimental datasets covering mouse and human cells using four different experiment protocols (Sec. 3.3 and 3.4). Our results show that existing methods give poor gene co-expression estimations and tend to have high false positive rates, thus, suggesting that robust estimation methods will need to focus on lowering the false positive rates. We also find, as suspected, that high sparsity in scRNA-seq data is the main bottleneck of co-expression estimation (Sec. 3.5). However, our implementation of several scRNA-seq data pre-processing approaches, including normalization (Stuart et al., 2019; Wolf et al., 2018; Choudhary and Satija, 2022; Borella et al., 2022), pseudo-bulk estimation (Squair et al., 2021), and imputation (Hou et al., 2020), show that while they reduce the sparsity and variance of the data, these pre-processing steps worsen the gene co-expression estimation by inducing biases (Sec. 3.6), for example, the imputation inducing some artificial correlations (Andrews and Hemberg, 2018; Hou et al., 2020). Therefore, we suggest that future works carefully choose pre-processing approaches when performing downstream estimation tasks from single-cell datasets. Previous works (Song et al., 2012; Ovens et al., 2021; Greenfield et al., 2010) tested several co-expression estimators, but they used microarray and bulk RNA-seq data. (Pratapa et al., 2020; Dibaeinia and Sinha, 2020) used scRNA-seq data in their benchmarks and found existing methods are largely inaccurate, but they focused on GRN estimation methods and did not examine the effect of different pre-processing steps. But our study gives a comprehensive investigation of co-expression estimations and is a novel supplement and extension of the earlier studies. Overall, as gene co-expression estimation is a fundamental step for downstream single-cell data analysis, our study provides valuable insights for this community that incorporate this step into their research.

## 2 Methods

### Notations

We let bold lowercase letters denote vectors, and bold uppercase letters denote matrices. For a matrix **A** ∈ ℝ*m*×*n*, its *i*-th row and *j*-th column are denoted by **A**_*i*:_ and **A**_:*j*_ respectively. We let 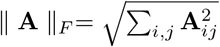 denote the Frobenius norm of matrix **A**. Given a vector **a** ∈ ℝ*n*, We let 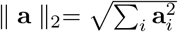 denote its ℓ_2_ norm.

We conduct a series of experiments to compare gene co-expression estimation methods on both simulated and real-world experimental scRNA-seq datasets. We first show that previously used simulations do not exhibit properties of real-world experimental data and use two recent single-cell read count simulation algorithms to generate realistic single-cell datasets (Sec. 3.1 and 3.2). Next, we compare nine bulk and single-cell gene co-expression estimators on our simulated and real-world datasets (Sec. 3.3 and 3.4). Finally, we investigate whether commonly used pre-processing approaches for scRNA-seq data could help extract robust gene co-expression networks (Sec. 3.6). We briefly introduce our experimental settings in this section (with details in the **Appendix: More on Method and Results** section), and Fig. 2 gives an overview of our setup.

**Figure 2:**
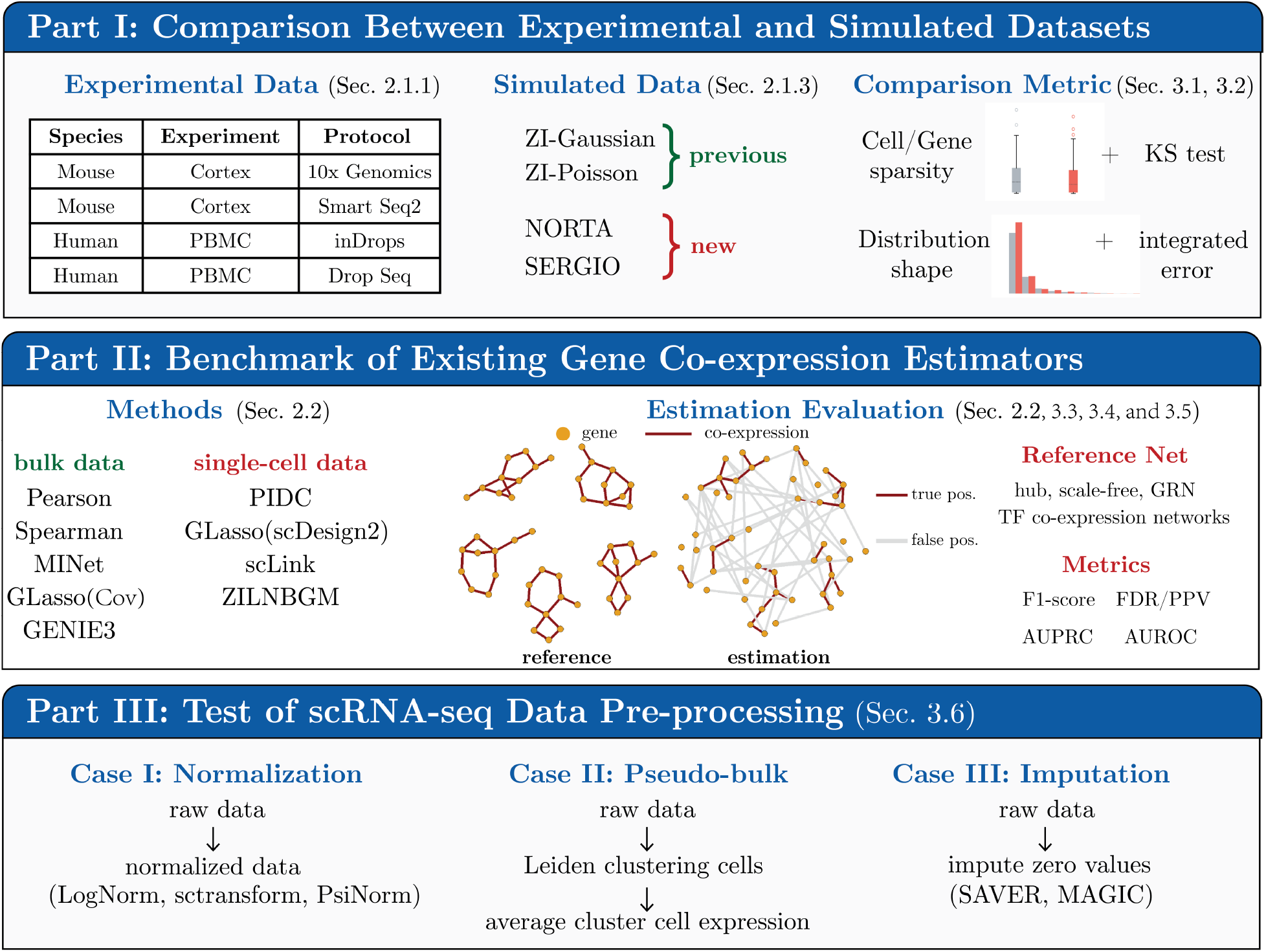
The summary of our study. **Part I**: We compare two previous and two new simulated processes with eight experimental data of multiple species and protocols in Sec. 3.1 and 3.2. We compare cell/gene sparsity and distribution shape qualitatively using plots and quantitatively using two-sample Kolmogorov–Smirnov tests (KS test) and integrated errors. **Part II**: We benchmark nine gene co-expression estimators, including both bulk and single-cell models, on simulated and experimental data in Sec. 3.3 and 3.4. We test three graph structures for simulation data: hub-based, scale-free, and gene regulatory network (GRN). We take known transcription factor (TF) co-expression networks for experimental data as reference networks. We report F1-score, FDR, AUROC, and AUPRC for method performances. **Part III**: We test multiple pre-processing approaches, including normalization, pseudo-bulk, and imputation on scRNA-seq data in Sec. 3.6.

### 2.1 Data Preparation

#### 2.1.1 Experimental Data and Pre-Processing Steps

We use eight publicly available scRNA-seq datasets in this study. We extract data of four different sequencing protocols from mouse cortex data (Ding et al., 2019) and human peripheral blood mononuclear cells (PBMC) datasets (Ding et al., 2019) downloaded from Single Cell Protal^3^. We filter out cells and genes with zero counts from each dataset’s raw unique molecular identifier (UMI) count matrix. The basic statistics of these eight datasets are summarized in Table 1. These datasets cover various protocols, cell library sizes, sparsity, and species. In our study, we select the top 100 and 1000 most highly variable genes (HVG) of every experimental dataset, and we conduct experiments on these subsets of data and corresponding simulations. The sparsity of 100 HVGs of each dataset are provided in Appendix Table A2.

**Table 1:** Basic statistics of eight experimental datasets.

| ID | Species | Experiment | Protocol | # Cells | # Genes | Sparsity<br>(ratio of non-zeros) | Max Count | Description |
| --- | --- | --- | --- | --- | --- | --- | --- | --- |
| #1 | Mouse | Cortex 1 | 10x Genomics | 1453 | 21887 | 10.95% | 4568 | Mouse cortex data. |
| #2 | Mouse | Cortex 1 | Smart Seq2 | 295 | 24114 | 22.94% | 108522 |  |
| #3 | Mouse | Cortex 2 | 10x Genomics | 4267 | 22805 | 7.58% | 3268 |  |
| #4 | Mouse | Cortex 2 | Smart Seq2 | 349 | 23974 | 20.48% | 59321 |  |
| #5 | Human | PBMC 1 | Drop Seq | 10824 | 22792 | 2.19% | 316 | Human PBMC. |
| #6 | Human | PBMC 1 | inDrops | 6184 | 18964 | 1.92% | 110 |  |
| #7 | Human | PBMC 2 | Drop Seq | 11578 | 24634 | 3.29% | 2623 |  |
| #8 | Human | PBMC 2 | inDrops | 5166 | 19648 | 3.19% | 553 |  |

#### 2.1.2 Gene Co-expression Reference Network Generation

To validate the performance of gene co-expression estimators, we need to generate gene co-expression networks to serve as the ground truth and simulate gene expression data based on it. In literature, hub genes (i.e., genes with high connectivity) are seen as highly essential and considered to play important roles in developmental processes and the regulation of transcription (Tong et al., 2004; Liu et al., 2019; Seo et al., 2009). Therefore, we generated hub-based networks as true co-expression network structures, denoted as **G**\* ∈ {−1, 0, 1} ^*p*×*p*^ with 1 and -1 indicating positive and negative edge respectively, in all simulations. We use `pcorSimulator` R package to generate networks. Fig. 8 gives an example of the hub-based network. For each network **G**\*, we obtain the network weight matrix **W**\* ∈ ℝ*p*×*p* through the following steps:

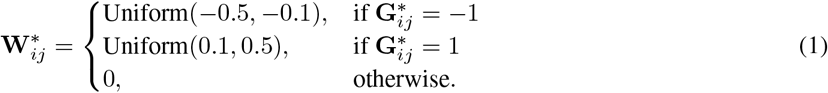

allowing us to use **W**\* in data simulation (see Sec. 2.1.3).

To define the ground truth networks for real-world experimental datasets (Sec. 3.4), we use known gene-gene interaction networks as references of gene co-expression (i.e., **W**\*). We focus on co-expression related to known transcription factors (TFs). We take the DNA-binding TF co-expression network presented in Zhou et al. (2017) as the reference for mouse datasets. Because it is constructed based on Pearson correlation, we keep edges with their absolute Pearson correlation coefficients higher than 0.2 to remove lowly co-expressed edges. For human datasets, we use the gene co-expression discovered in ChIP-seq experiments Rouillard et al. (2016). We use the gene cosine similarity matrix constructed from Pathway Common Rodchenkov et al. (2020) dataset, and select the sub-network related to human TFs Lambert et al. (2018). We keep the edges with their absolute cosine similarity higher than 0.2. The average human/mouse TF co-expression network sparsity (i.e., the number of edges divided by the number of possible gene pairs) is 11%.

#### 2.1.3 scRNA-seq Data Simulation

Previous studies have used two representative simulations based on Gaussian and Poisson distributions to evaluate state-of-the-art gene co-expression estimation methods.

- **Zero-inflated Gaussian (**`ZI-Gaussian`): The Gaussian-based simulation strategy in (Li and Li, 2021; Mestres et al., 2018) assumes non-zero values of the normalized gene expression matrix following a Gaussian distribution. This strategy first generates a co-expression network **G** following Sec. 2.1.2 and constructs a partial correlation matrix (i.e., the inverse of the covariance matrix) **Σ**^−1^ where 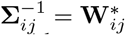 denotes the edge weight between node *i* and *j*. Then it uses this partial correlation matrix along with gene expression means and standard deviations estimated from real scRNA-seq data to simulate a dense gene expression matrix 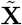 through a multivariate Gaussian distribution. To introduce sparsity into the matrix, for each matrix element, it calculates the probability 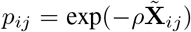 where *ρ* is a hyper-parameter controlling the level of sparsity. The final sparse simulation matrix is obtained by

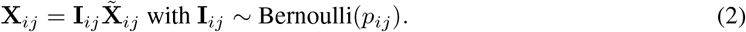

Following the same experiment setting in (Li and Li, 2021), we simulated four datasets with the same partial correlation matrix and different *ρ* ∈ {0.07, 0.10, 0.13, 0.16} in our study. Since this simulation adds zeros on top of Gaussian samples, we name it `ZI-Gaussian` in our study.

- **Zero-inflated Poisson (**`ZI-Poisson`): In addition, (Choi et al., 2017; Allen and Liu, 2013) use Poisson-based strategy. It first generates a gene expression matrix 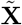 through a linear combination

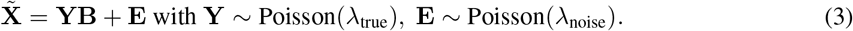

Here, **B** is a permutation matrix based on **W**\*. In order to have zeros, it then multiplies each element in the 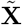 with a Bernoulli random variable as

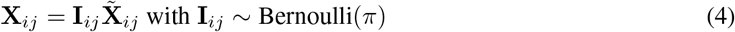

where *π* controls the data sparsity. Detailed simulation process may refer to (Choi et al., 2017). We named this Poisson-based simulation as `ZI-Poisson`. Adopting the same experiment setting in (Choi et al., 2017), we generated three simulation datasets by fixing *λ*_true_ = 1.5, *λ*_noise_ = 0.5, and varying *π* ∈ *{*0.8, 0.9, 1.0*}*.

Because `ZI-Gaussian` and `ZI-Poisson` simulations cannot exhibit characteristics of experimental scRNA-seq data (see Sec. 3.1), we apply two new simulation processes.

- **NORmal-To-Anything (**`NORTA`): The first one is `NORTA` based on the normal-to-anything approach (Cario and Nelson, 1997; Chen, 2001; Sekula et al., 2020). The essential idea of `NORTA` is transforming multivariate Gaussian samples to samples with any given marginal distributions while preserving a given covariance. Previously, Sun *et al*. (Sun et al., 2021) have shown that using information from experimental data helps with mimicking the properties of real single-cell data. So in our study, we use `NORTA` to curate data with distributions learned from experimental data. Concretely, given experimental scRNA-seq data of *p* genes, we first use Negative Binomial (NB) distributions to fit each gene’s expression because NB is known to fit scRNA-seq data well (Hafemeister and Satija, 2019; Svensson, 2020), and obtain *p* marginal distributions. Then, we generate the covariance matrix as **Σ** = **W**\*. `NORTA` utilizes them to simulate data with these *p* marginal NB distributions and gene covariance encoded by **Σ**. Fig. 3 illustrates `NORTA` simulation. For every experimental dataset, we generate corresponding synthetic data with `NORTA` ^4^. In `NORTA` simulations, the average hub-based network sparsity is 2.3%.
- **Single-cell ExpRession of Genes In silicO (**`SERGIO`): Another new simulation strategy is `SERGIO` (Dibaeinia and Sinha, 2020). Given a GRN and profiles of regulators, `SERGIO` models the stochasticity of transcriptions and regulators with stochastic differential equations (SDEs). Concretely, it first generates dense gene expression matrix 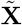 in logarithmic scale at stationary state via SDEs. Then it adds zeros via

**Figure 3:**
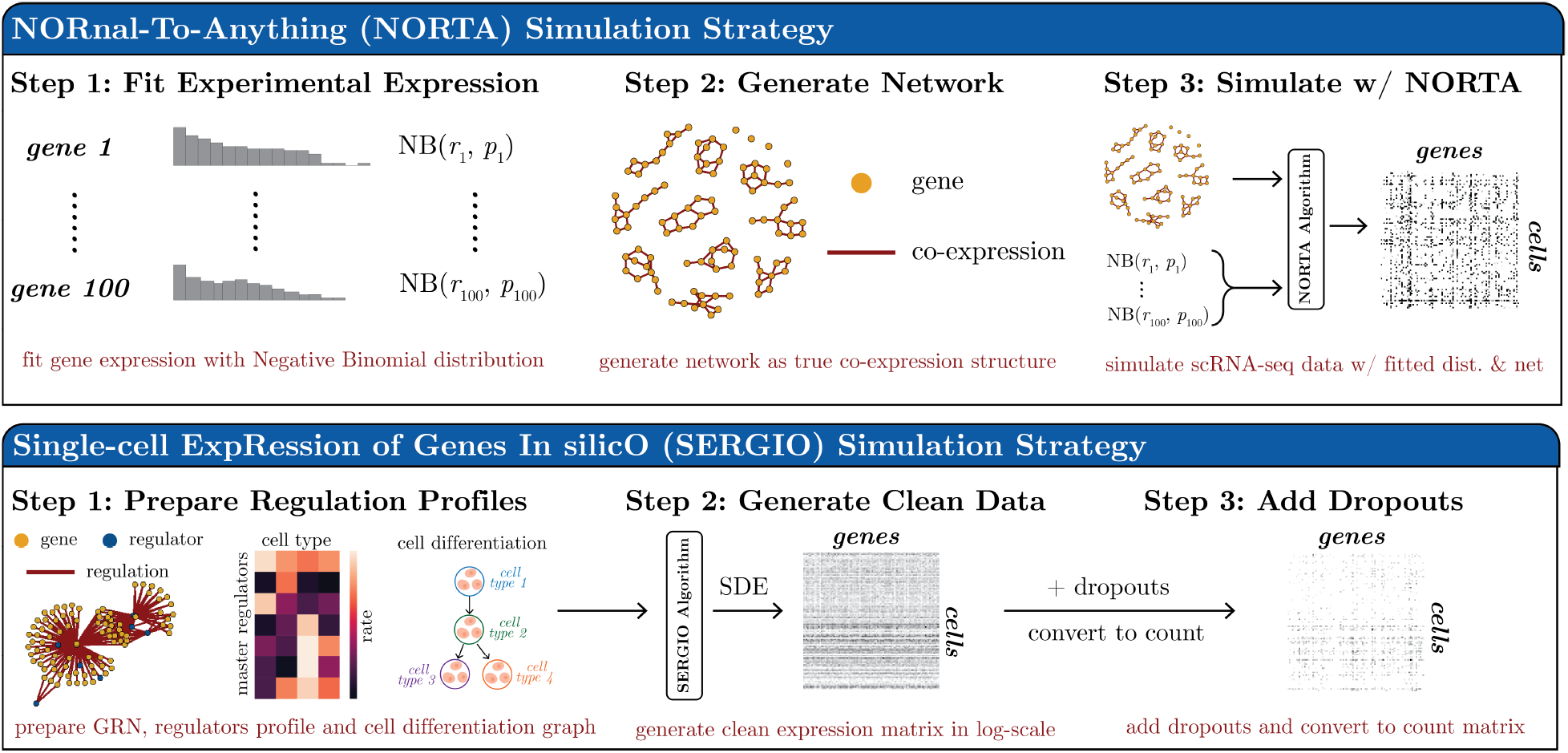
Illustration of `NORTA` and `SERGIO` simulations. **(Top)** `NORTA` simulation strategy. First, it fits experimental data gene expression with negative binomial distributions. It generates a network encoding the covariance matrix. Then, the fitted negative binomial distributions and covariance matrix are used to simulate scRNA-seq data. **(Bottom)** `SERGIO` simulation strategy. It first prepares GRN, regulators profile (i.e., basal production rate in each cell type), and cell type differentiation graph. Then using these data, it generates clean data based on stochastic differential equations (SDEs). Finally, it adds dropouts to the clean expression matrix and converts it to the count expression matrix.

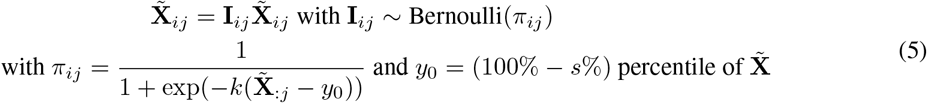

with two hyper-parameters *k* and *s* determining the sparsity. Finally, it convert 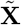 to the expression matrix through

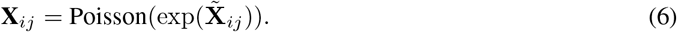

The paper (Dibaeinia and Sinha, 2020) provides a comprehensive experiment showing that `SERGIO` can simulate realistic scRNA-seq data. Fig. 3 provides an illustration of `SERGIO` simulation. In our study, we simulate five datasets with *k* = 6.5 and *s* = 1, 5, 10, 15, 20 using the GRN and regulator profiles provided by (Dibaeinia and Sinha, 2020). In `SERGIO` simulations, the GRN sparsity is 5.2%.

In `ZI-Gaussian` and `ZI-Poisson` simulations, we set the number of cells as 1000. In `SERGIO` simulation, the number of cells produced is 2700, with nine cell types in total and 300 cells per cell type. For `NORTA`, the number of cells in simulations equals that in the corresponding experimental data. So in all cases, the number of cells is much larger than the number of genes. We compare the characteristics of these simulation datasets with experimental datasets in Sec. 3.1,3.2 and test methods on simulations in Sec. 3.3.

### 2.2 Gene Co-Expression Network Estimators and Evaluation Metrics

We have tested nine gene co-expression network estimation methods in this study. These methods are summarized in Table 2. They consist of five methods designed for microarray or bulk RNA-seq data and four methods for scRNA-seq data. These methods cover a wide range of state-of-the-art and commonly used gene co-expression estimators with different data distribution assumptions and co-expression measurements. Detailed descriptions are provided in **Appendix: More on Method and Results** section.

**Table 2:** Summary of gene co-expression estimation methods tested in this study. The first five are bulk data methods and the rest four are single-cell data methods. “NA” means the method has no explicit assumption on the data distribution.

| Name | Citation | Co-expression Measurement | Network Estimation | Hyper-parameter | Assumption |
| --- | --- | --- | --- | --- | --- |
| Pearson | (Langfelder and Horvath, 2008) | Pearson correlation | thresholding | threshold $\delta$ | NA |
| Spearman | (Langfelder and Horvath, 2008) | Spearman correlation | thresholding | threshold $\delta$ | NA |
| MINet | (Meyer et al., 2008) | mutual information | thresholding | threshold $\delta$ | NA |
| GLasso(Cov) | (Friedman et al., 2008) | sample covariance | MLE | regularization $\lambda$ | Gaussian |
| GENIE3 | (Huynh-Thu et al., 2010) | regression trees | thresholding | threshold $\delta$ | NA |
| PIDC | (Langfelder and Horvath, 2008) | partial information | thresholding | threshold $\delta$ | gamma or Gaussian |
| scLink | (Li and Li, 2021) | robust correlation | MLE | regularization $\lambda$ | gamma-Gaussian mixture |
| GLasso(scDesign2) | (Sun et al., 2021) | copula correlation | MLE | regularization $\lambda$ | zero-inflated negative binomial |
| ZILNBGM | (Park et al., 2021) | regression coefficient | regression | regularization $\lambda$ | zero-inflated negative binomial |

To evaluate the performance of methods, we consider co-expression estimation a binary classification task and compare estimated co-expression structures with the reference structure. Specifically, let **G**\* ∈ {0, 1} ^*p*×*p*^ denote the true co-expression of *p* genes ^5^ with 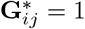 representing genes *i* and *j* are co-expressed. So all 1s in **G** are considered positives, and 0s are negatives. 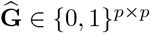 denotes the estimated co-expression. We evaluate methods with these metrics:

#### F1-score

The F1-score of estimation is computed with the upper triangle (without diagonal entries) of ground truth **G**\* and prediction 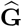, because the co-expression matrix is a symmetric matrix with diagonal entries all 1s:

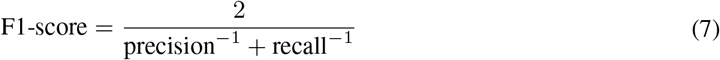

with

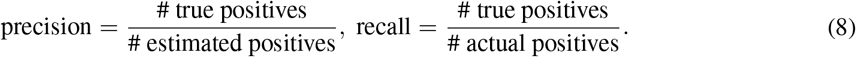

#### Ratio of false positives

To evaluate how much false positives each method produce, we computed false discovery rate (FDR) and positive predictive value (PPV), or precision, with

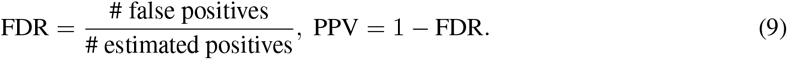

All the tested methods have only one hyper-parameter to control the sparsity of estimated networks. Specifically, `Pearson`, `Spearman`, `MINet`, `GENIE3`, and `PIDC` require a threshold *δ* to filter small values. So in our experiments, we generated a sequence of 100 equally spaced *δ* values from 0 to the maximal value in the measurement matrix to obtain a sequence of estimations from the complete network to the empty network. Then the final estimation is selected based on the highest F1-score. On the other hand, `GLasso(Cov)`, `GLasso(scDesign2)`, `scLink`, and `ZILNBGM` have *λ* as the sparsity-regularization hyper-parameter. Similarly, we generated a sequence of 100 equally spaced *λ* values from 0 to 1 and picked the best estimation with the highest F1-score. Especially, `ZILNBGM` suggests the maximal *λ* as the max covariance value. So we used this setting only for `ZILNBGM`. We also include the area under the receiver operating characteristic curve (**AUROC**) and the area under the precision-recall curve (**AUPRC**) values. For each of the 100 trials, we compute the estimation’s precision, recall, true positive rates, and false positive rates. AUPRC and AUROC are calculated based on curves constructed from these 100 metric values.

## 3 Results

### 3.1 Existing simulations to test co-expression estimation methods do not capture real-world data properties

We first test if two previously-used simulation processes, `ZI-Gaussian` and `ZI-Poisson`, represent real-world data. We simulate ‘single-cell’ datasets with 100 genes and hub-based gene co-expression networks, following the parameter setting in (Li and Li, 2021; Choi et al., 2017). We compare the properties of these simulations with experimental datasets from Table 1. Note that for experimental data, we take 100 HVGs from tens of thousands of genes; while on simulations, we directly generate 100 genes without a selection step. The HVG selection takes informative genes, and simulated genes are naturally informative (since we are not simulating uninformative noise), so the experimental and simulated data here (and in Sec 3.2) are comparable.

We systematically compare the sparsity (i.e., the ratio of non-zero values over total values) per cell and sparsity per gene between 100 HVGs of eight experimental datasets and seven `ZI-Gaussian` and `ZI-Poisson` simulated datasets with 100 genes. Fig. 4(A-B) show simulations of both processes contain much fewer zeros than the experimental data. The average sparsity per cell for all eight experimental data is 0.083. But the most sparse `ZI-Gaussian` simulated dataset (i.e., *ρ* = 0.07) has average sparsity per cell of 0.468; the most sparse `ZI-Poisson` simulated dataset (i.e., *π* = 0.8) has average sparsity per cell of 0.789. Similarly, the average gene sparsity of experimental data is 0.083, while it of the most sparse `ZI-Gaussian` simulation is 0.47 and the most sparse `ZI-Poisson` simulation is 0.79. Two-sample Kolmogorov–Smirnov tests (KS test) between every experimental data and every simulated data give *p <* 10^−14^ (for cell sparsity) and *p <* 10^−16^ (for gene sparsity). This demonstrates that the cell and gene sparsity of simulated data is significantly different from experimental data.

**Figure 4:**
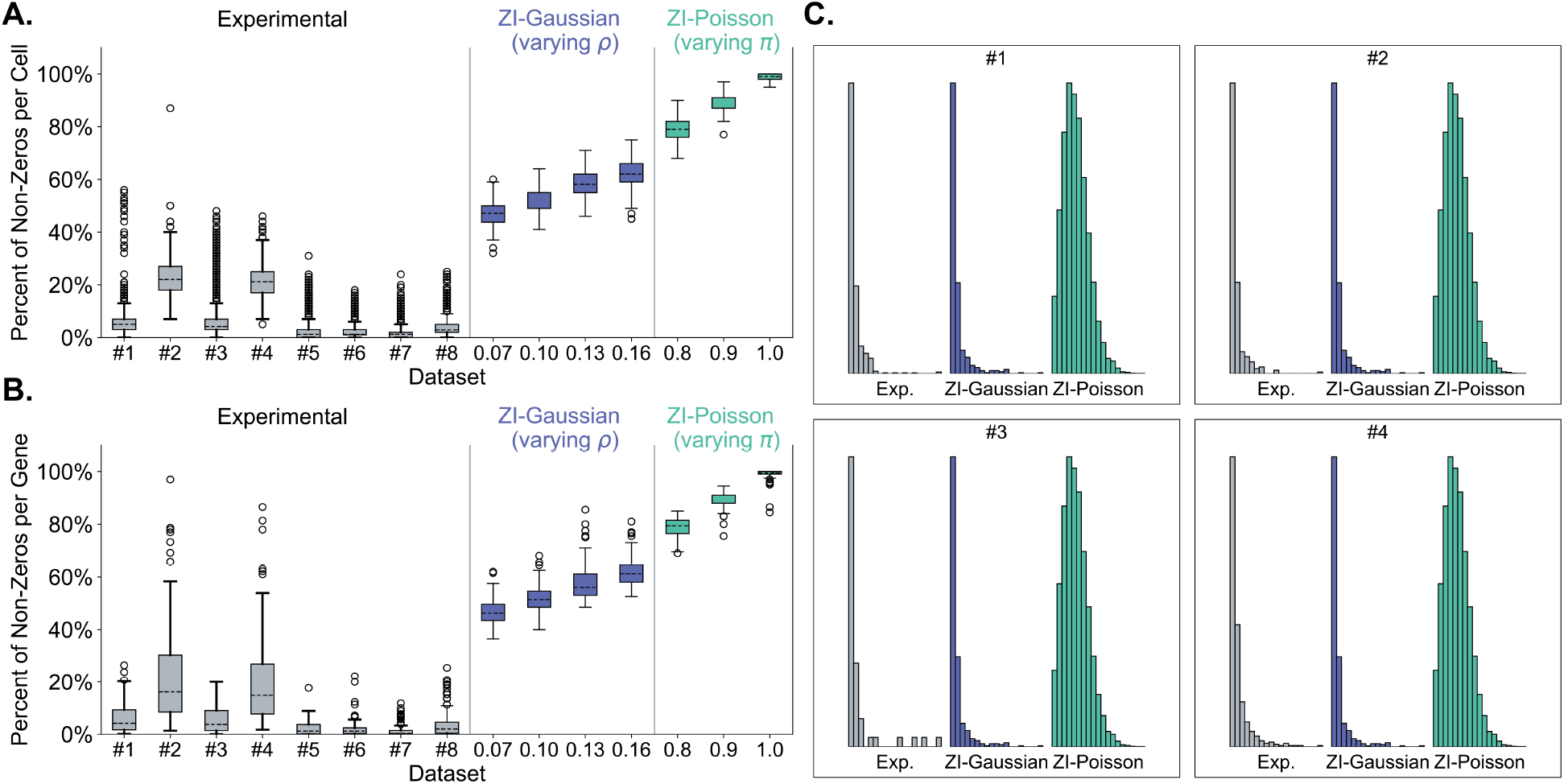
Compare properties between experimental data, `ZI-Gaussian` (with *ρ* = {0.07, 0.10, 0.13, 0.16}), and `ZI-Poisson` (with *π* = {0.8, 0.9, 1.0}) simulations. (**A**) Sparsity per cell. (**B**) Sparsity per gene. (**C**) Compare four experimental datasets (#1 to #4) with the most sparse `ZI-Gaussian`/`ZI-Poisson` simulations. Histograms show distributions of non-zero parts for min-max scaled data.

Furthermore, we compare the expression distribution shape. Because the value range differs among datasets, we first normalize each dataset by min-max scaling as

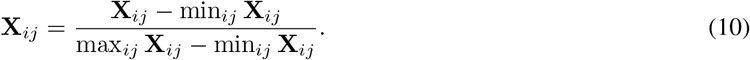

Data distributions of all datasets have a bi-modal structure consisting of dominating zero and rest non-zero parts. Because datasets have different sparsity levels, we only compare the distribution of non-zero components in Fig. 4(C). `ZI-Poisson` simulations are qualitatively different from experimental data. Also, we compute integrated absolute error (IAE) as

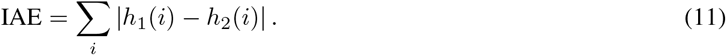

Here *h*_1_ and *h*_2_ are 20-bins histograms of two datasets, and *h*_1_(*i*) is the data frequency at the *i*-th bin. The average IAE between `ZI-Gaussian` and experimental data is 0.04; between `ZI-Poisson` and experimental data is 1.73; between every possible pair of experimental data is 0.03. This further indicates `ZI-Poisson` simulations are different from experimental data.

Our results show that `ZI-Gaussian` and `ZI-Poisson` simulations cannot display properties of actual scRNA-seq data. They are either less sparse or have very different distribution shapes. Therefore, despite good performance, the reliability of methods tested on these simulations becomes less certain. Next, we perform a comprehensive benchmark of gene co-expression estimation methods by generating more realistic simulations.

### 3.2 Generating Simulation with Experimental Data properties to Help Validate co-expression estimation Performance

We apply two new simulation processes, namely `SERGIO` and `NORTA`, to generate more realistic data. To validate that the two new simulation processes can represent real-world experimental data, we also compare cell sparsity, gene sparsity, and distribution shapes of the normalized data between these new simulations and experimental datasets. Fig. 5(A-B) and Appendix Fig. A1 qualitatively show that `NORTA` and `SERGIO` simulation datasets have the similar gene and cell sparsity as the experimental data. The average gene sparsity of `NORTA` simulations is 0.087, and of `SERGIO` simulations is 0.11, which is similar to the average gene sparsity of experimental data (0.083). The cell sparsity of `NORTA` simulations is 0.087, and of `SERGIO` simulations is 0.11, which is also similar to the experimental data average cell sparsity (0.083). Two-sample KS test between most `NORTA` simulation and corresponding experimental data gene sparsity gives *p >* 0.1, except for dataset #6 (*p* = 1.2 × 10^−4^) and #8 (*p* = 2.4 × 10^−2^). Because these two experimental datasets have genes with very high sparsity levels, e.g., 0.03% non-zeros, the `NORTA` generated all zeros for these genes. So the KS test gave p-values smaller than 0.1 in these cases, but overall the `NORTA` can accurately simulate the sparsity level of the experimental data. KS tests between `SERGIO` simulation and experimental data gene sparsity all give *p >* 0.1. These denote no significant differences between simulation gene sparsity and experimental data gene sparsity. Moreover, Fig. 5(C-D) show that new simulations have similar distribution shapes with experimental data, with `NORTA` ISE=0.014 and `SERGIO` ISE=0.086. Our results demonstrate that `SERGIO` and `NORTA` better exhibit the properties of the experimental data compared to the previous simulation methods.

**Figure 5:**
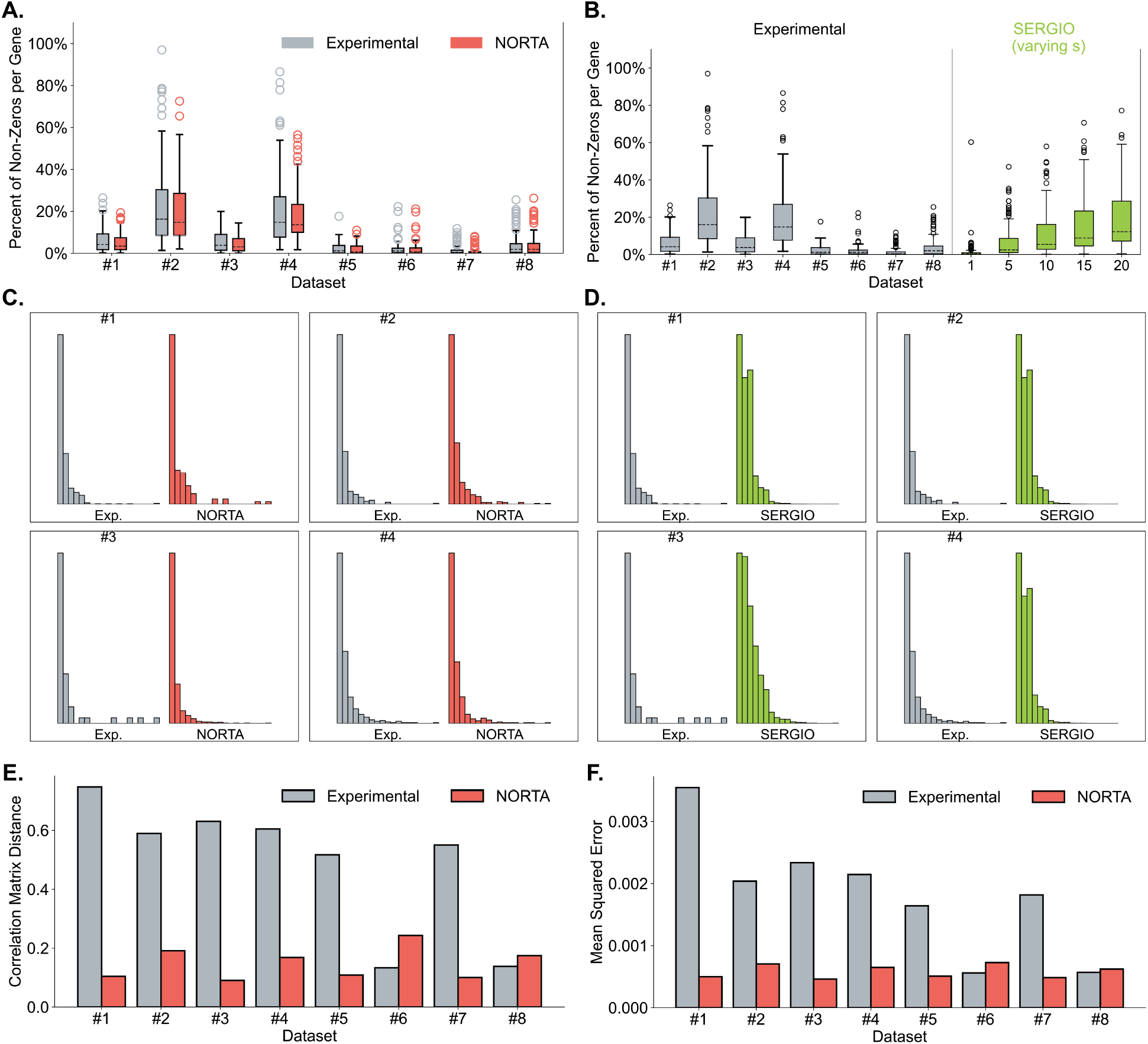
Comparison between experimental data, `NORTA`, and `SERGIO` simulations. (**A**) Sparsity per gene of experimental data and corresponding `NORTA` simulations. (**B**) Sparsity per gene of experimental data and `SERGIO` simulations. (**C**) Compare distribution shapes between four experimental datasets and corresponding four `NORTA` simulations. Histograms show distributions of non-zero parts for min-max scaled data. (**D**) Compare distribution shapes between four experimental datasets and `SERGIO` simulations. Histograms show distributions of non-zero parts for min-max scaled data. (**E**) Correlation matrix distance (CMD) of experimental data and corresponding `NORTA` simulations. The gray bar denotes CMD between experimental data correlation and true network, i.e., CMD(Corr_exp_, **W**\*). The red bars denotes CMD between `NORTA` simulation correlation and true network, i.e., CMD(Corr_sim_, **W**\*). (**F**) Mean squared error (MSE) of experimental data and corresponding `NORTA` simulations. The gray bar denotes MSE between experimental data correlation and true network, i.e., MSE(Corr_exp_, **W**\*). The red bar denotes the MSE between `NORTA` simulation correlation and true network, i.e., MSE(Corr_sim_, **W**\*).

Note that experimental single-cell data generally contains gene expressions of multiples cell types (Consortium et al., 2018; Han et al., 2018; Rozenblatt-Rosen et al., 2017), so we check whether the `NORTA` and `SERGIO` simulations also have such heterogeneous information via clustering. We find the first 50 principal components for each dataset through the Principal Component Analysis (PCA), and we cluster cells with the Leiden algorithm (Traag et al., 2019). Leiden algorithm is a community detection method used in many scRNA-seq clustering algorithms (Cao et al., 2019; Mohammadi et al., 2020; Stuart et al., 2019; Wolf et al., 2018) that can automatically choose the number of clusters and show superior performance on finding cell types from single-cell data (Yu et al., 2022). We visualize each dataset’s clustering result in the 2D Uniform Manifold Approximation and Projection (UMAP) (McInnes et al., 2018) embedding space. Quantitatively, we compute the silhouette score (Rousseeuw, 1987) in the UMAP space to evaluate the cohesion and separation of clusters. The silhouette score ranges from -1 to +1, with a higher value indicating dense and well-separated clusters and values around zero indicating overlapping clusters. Fig. 6 shows that the `NORTA` and `SERGIO` simulations have clusters that are somewhat well separated (Appendix Fig. A2, A3, and A4 provide clustering visualization for all the `NORTA` and `SERGIO` simulations). Also, the `NORTA` simulation has a similar silhouette score with its corresponding experimental data. Note that we are not matching the cell clusters between experimental and simulated data. Having a silhouette score higher than 0 and similar to that on experimental data is sufficient to demonstrate that simulations possess multi-cell-type properties and are realistic.

**Figure 6:**
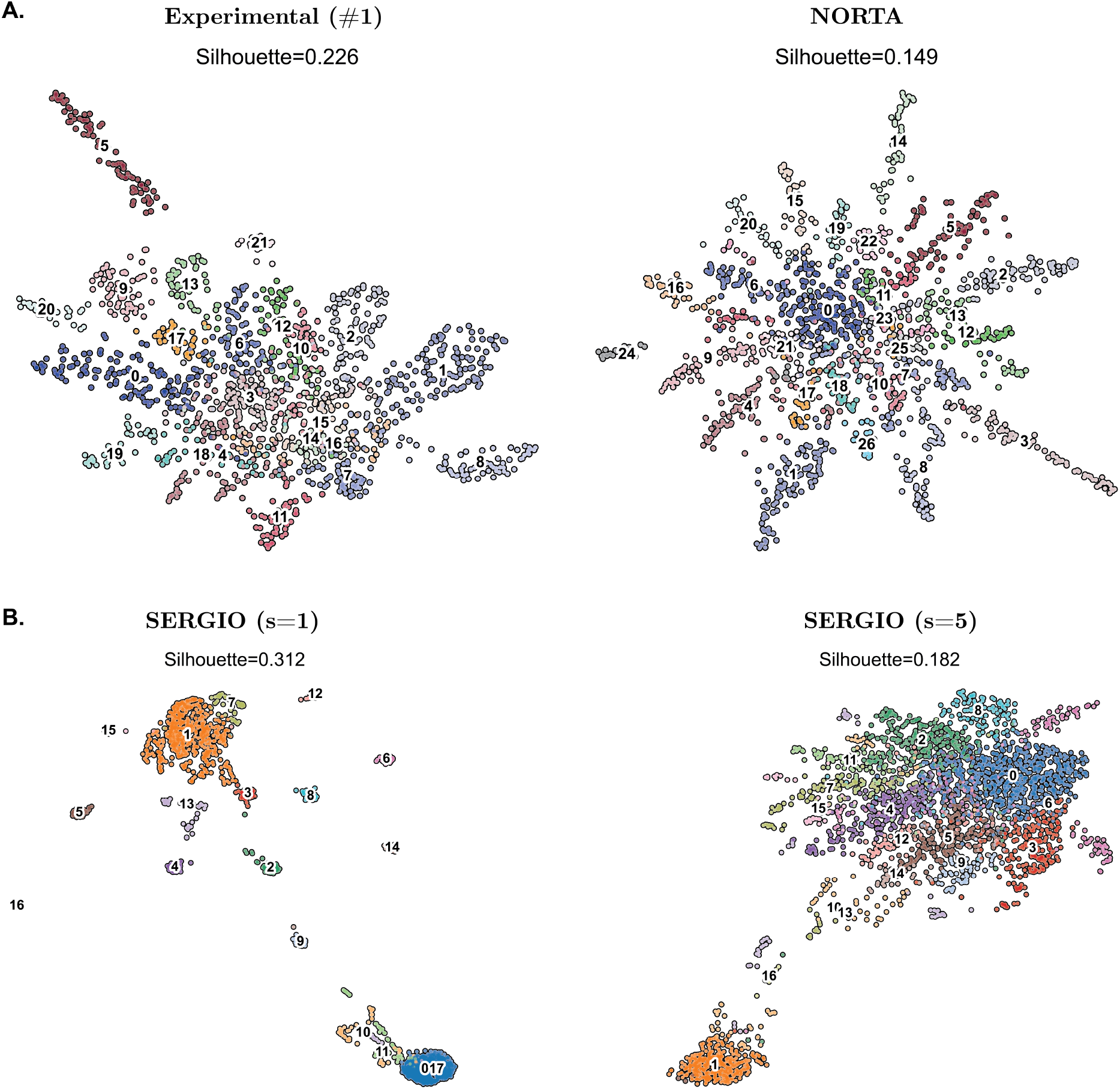
Examples of cell clustering on experimental data, `NORTA` simulations, and `SERGIO` simulations. UMAP embedding visualization for the Leiden clustering on the first 50 principal components. Silhouette score is computed in the UMAP space. (**A**.) Experimental dataset (#1) and its corresponding `NORTA` simulation. (**B**.) `SERGIO` simulations with simulation parameter *s* = 1 and *s* = 5.

Another important aspect of the simulation is imposing a given gene co-expression structure on data such that it is feasible for estimation. (Dibaeinia and Sinha, 2020) gave a comprehensive testing of `SERGIO` and found simulations successfully have specified co-expression patterns. So here, we only validate that `NORTA` simulations have information of specified gene co-expression by comparing the Pearson correlation coefficients between experimental data and corresponding simulations. Specifically, let Corr_exp_ denote Pearson correlation coefficient matrix of experimental data; Corr_sim_ denote correlation matrix of corresponding `NORTA` simulations. We use mean squared error (MSE) and correlation distance matrix (CMD) (Herdin et al., 2005) to measure the similarity between two correlation matrices. Here, a smaller MSE between two correlation matrices indicates they are more similar. CMD(**A, B**) = 0 denotes two correlation matrices are equal up to a scaling factor and CMD(**A, B**) = 1 if they differ in max extents. CMD values in Fig. 5(E-F) show that CMD(Corr_sim_, **W**\*) *<* CMD(Corr_exp_, **W**\*) and MSE(Corr_sim_, **W**\*) *<* MSE(Corr_exp_, **W**\*) in most cases. Exceptions are still datasets #6 and #8 for the reason mentioned above, having slightly higher CMD and MSE in simulations. But overall, `NORTA` successfully imposes the co-expression structure **W**\* on data.

In summary, the `SERGIO` and `NORTA` are more realistic simulations and better options in future works for evaluating gene co-expression estimators.

### 3.3 Existing Methods Estimate Gene Co-Expression Networks from realistically Simulated Data with a high FDR

Here, we test nine state-of-the-art gene co-expression estimators on `SERGIO` and `NORTA` simulations. We compute and compare F1-score, FDR, AUROC, and AUPRC across all methods.

On `NORTA` simulated datasets, some methods including the state-of-the-art scRNA-seq model `scLink`, can have incorrect predictions with F1-score lower than 0.2 (Fig. 7(A)). In other cases, even though the F1-score is around 0.5 for `Spearman`, `MINet`, and `GLasso(scDesign2)`, they have FDR around 0.4 to 0.5 (Fig. 7(B)), indicating the predictions have many false positives. Appendix Fig. A11 gives an example of `Spearman` method prediction on `NORTA` simulations with 100 genes. In this case, F1-score=0.43 and FDR=0.6, containing 27 true positives and 40 false positives. This denotes that although F1-score is relatively high, the predictions have many false positives. On `SERGIO` simulations, all methods have F1-score lower than 0.2 and FDR higher than 0.8, giving poor co-expression estimations (Fig. 7(C-D)). But we still can use a few methods piratically. Suppose one has to select a co-expression estimator to give a coarse-grained estimation as the starting point of GRN construction. In that case, we recommend using `Spearman`, `MINet`, or `GLasso(scDesign2)` since they have better performance in our experiments. The reason for it is that Spearman correlation and mutual information are robust to outliers and noise, and `GLasso(scDesign2)` captures gene correlations with Gaussian copula (Sun et al., 2021).

**Figure 7:**
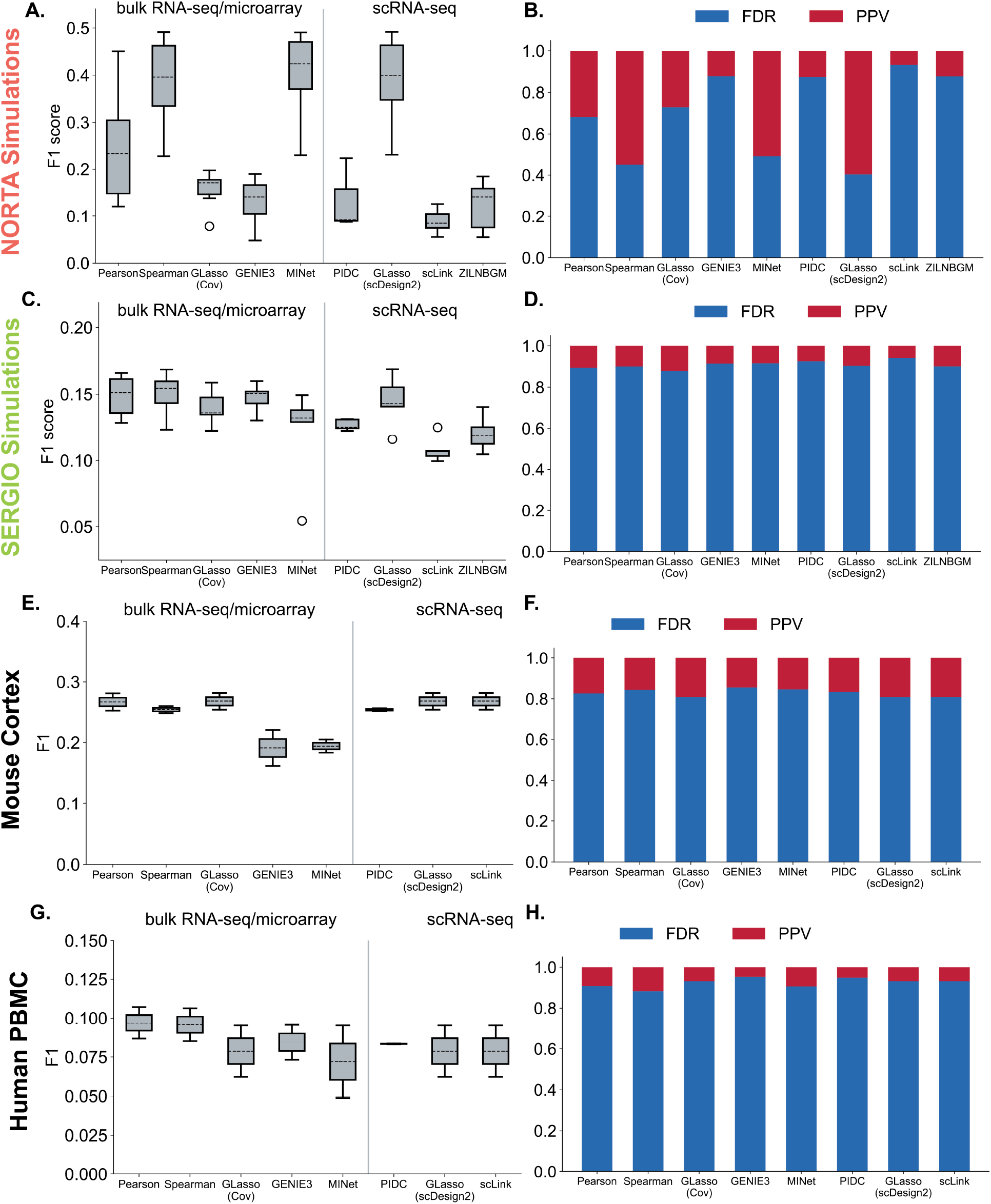
Methods performances on simulated and experimental datasets. In box plots, we split the performance of five bulk RNA/microarray models and four scRNA-seq models with a vertical line. Because `ZILNBGM` spends several days running a single trial on experimental data, we consider it uncomparable with other methods and omit it in the plots related to experimental data. (**A-B**) F1-score and FDR/PPV on `NORTA` simulations with 100 highly variable genes. (**C-D**) F1-score and FDR/PPV on `SERGIO` simulations with 100 genes. (**E-F**) F1-score and FDR/PPV on mouse cortex data with 1000 highly variable genes. (**G-H**) F1-score and FDR/PPV on human PBMC data with 1000 highly variable genes.

In Appendix Fig. A7 and A8, we tested methods on `NORTA` and `SERGIO` simulated data with more genes. It shows that methods’ performance decrease with an increased number of genes (see **Appendix Section A1.5**). This is reasonable since having more genes indicates the problem has higher complexity. So in later sections, we only use 100-genes simulations.

We also notice that methods have high AUROC but much lower AUPRC values (Appendix Fig. A5, A6). The reason for high AUROC is that the ground-truth network is very sparse, such that the number of negative instances (i.e., absence of co-expression) is much larger than the number of positive instances (i.e., the existence of co-expression). The AUROC measurement is known to be sensitive to class imbalance, so we also use AUPRC score, which alleviates the influence of data imbalance by considering both precision and recall values. Our observations suggest that co-expression estimation studies should use multiple metrics to comprehensively evaluate their methods.

Finally, we test methods on several simulated datasets from the previous studies for the completeness of comparison. First, we compare method estimations on `ZI-Gaussian` and `ZI-Poisson` simulations generated in Sec. 3.1. On four `ZI-Gaussian` simulations, `scLink` outperforms other models with average F1-score=0.29, average FDR=0.74, and average AUPRC=0.22 (Appendix Fig. A9). The high F1-score of `scLink` is reasonable since the generation of `ZI-Gaussian` follows the Gaussian assumptions of the `scLink` method. We use the same simulation used in (Li and Li, 2021) and `scLink` achieves a similar AUPRC value as Fig. 2(D) in (Li and Li, 2021). However, though `scLink` outperforms its counterparts, its predictions still report a high FDR. On the other hand, on three `ZI-Poisson` simulations, Appendix Fig. A10 demonstrates that most methods have relatively better performance on `ZI-Poisson` than on other tested simulations (including `ZI-Gaussian`, `NORTA`, and `SERGIO`) and experimental data. Specifically, `scLink` is the best with average F1-score=0.75, average FDR=0.21, and average AUPRC=0.78. We think these good estimations may be due to the low sparsity of `ZI-Poisson` simulations but are not feasible in real-world single-cell applications due to their inconsistency with experimental data, as shown in Fig. 4. Through these experiments, we find that existing methods either report high FDR or use less sparse simulations than real-world data, making them less applicable to experimental single-cell datasets. On the contrary, we conduct experiments on more realistic simulations with comprehensive evaluations, which will be more reliable for future testing and benchmarking.

### 3.4 Methods Recover Known Gene Interactions from Experimental scRNA-seq Data with a low precision

Here we test methods on four experimental datasets (#2, #4, #5, and #7 in Table 1) with 1000 HVGs ^6^ to evaluate how they perform on real scRNA-seq data. We compare model predictions with known TF co-expression networks discovered on bulk data (see Sec. 2.1.2). For each experimental dataset, the number of TFs and the corresponding number of known co-expression edges are listed in Appendix Table A1. Note that these TF co-expression networks are not absolute ground truth since they are obtained from bulk data instead of single cells. But they are good references for us to evaluate whether methods can recover known information. Results in Fig. 7(E-H) show that all methods have less than ideal performances on detecting known TF co-expression from experimental data. All methods have average F1-score ≈ 0.25 and average PPV*<* 0.2 for mouse cortex data. `GLasso(Cov)`, `GLasso(scDesign2)`, and `scLink` perform best with an average PPV of 0.191. On the human PBMC data, these methods’ average F1-score*<* 0.1 and average PPV*<* 0.12, where `Spearman` outperforms others, having an average PPV= 0.117.

With these tests on experimental datasets, we find that our tested methods have a low precision when recovering bulk data-based TF-TF edges via their co-expression estimation from single-cell datasets. We also note that evaluating co-expression on experimental data needs careful consideration. A missing edge in reference networks does not mean the pair of genes are not co-expressed. That edge/co-expression perhaps has not been verified by previous bulk data experimental studies. So the FDR is expected to be high, as the estimator may discover unknown gene co-expression from single-cell data. We consider an alternative metric like precision (i.e., PPV), see Eq. 8. The precision/PPV evaluates how many estimated positives are true positives. This metric can test models’ performance on recovering known gene co-expression. However, since precision=1-FDR, high FDR values lead to low precision, and all methods give precision*<* 0.5 in our experiments. Unfortunately, we generally have little information to validate if the false positives are newly discovered co-expression interactions or errors. Therefore, we suggest caution while choosing the evaluation metric to verify co-expression estimation for real-world datasets. Augmenting downstream validations, such as the quality of GRN inference from the co-expression matrix and the accuracy of gene enrichment analysis, might be useful. Furthermore, we continue the remaining experiments on our simulations since we know the underlying ground truth.

### 3.5 High Sparsity in scRNA-Seq Data Impedes the Correlation Computation leading to poor Co-Expression Estimation

We generate data with different settings to explore which property of scRNA-seq data is the bottleneck of co-expression estimation. Since `NORTA` is more flexible and convenient at controlling simulation properties, like the type of network topology, we explore `NORTA` simulation in this section.

First, we use `NORTA` to simulate data with 100 genes and {10, 50, 100, 150, 200} cells. We observe that using more cells helps with the estimation (Fig. 9(A)). For scRNA-seq data, as the sequencing techniques enable the profiling of more cells, we can have data with hundreds and thousands of cells. So the number of cells is not the bottleneck.

Some works state that real-world biological networks can be scale-free networks (Khanin and Wit, 2006; Clauset et al., 2009; Albert, 2005). Additionally, using known gene regulatory networks (GRNs) in simulation is a good way to incorporate prior knowledge. So we further use scale-free networks and example GRNs from (Dibaeinia and Sinha, 2020) in `NORTA` simulations with 100 genes and compare them with hub-based networks. Fig. 8 gives an example of these three types of networks. Fig. 9(B-C) shows the similar performance of models, denoting that the graph structure is not the important factor.

**Figure 8:**
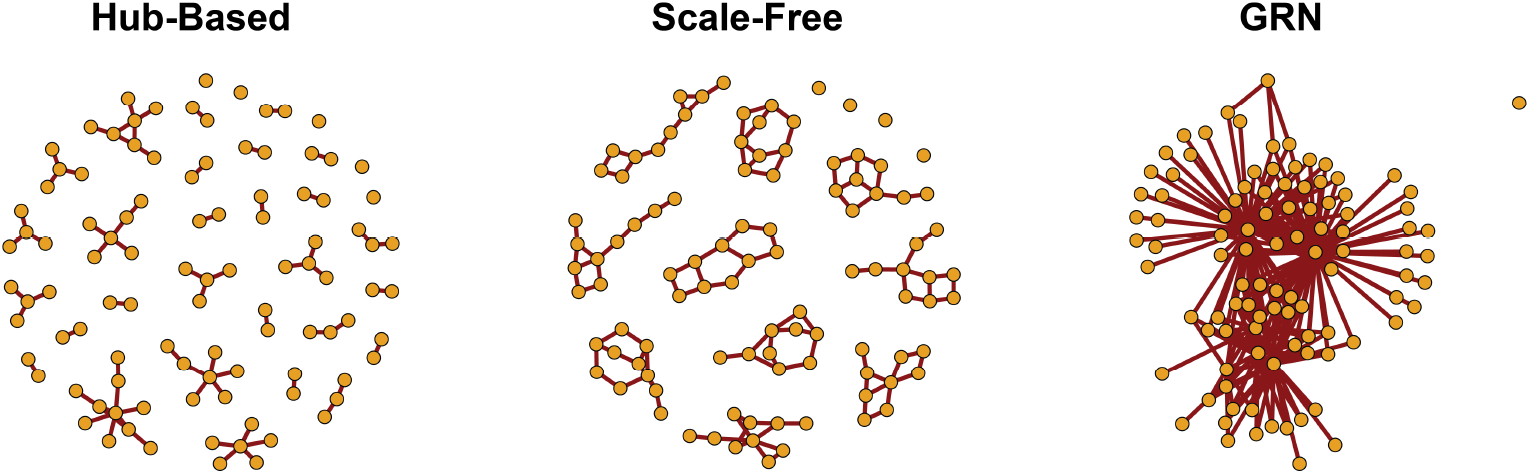
Examples of three gene co-expression network topologies used in simulations: hub-based network, scale-free network, and GRN.

**Figure 9:**
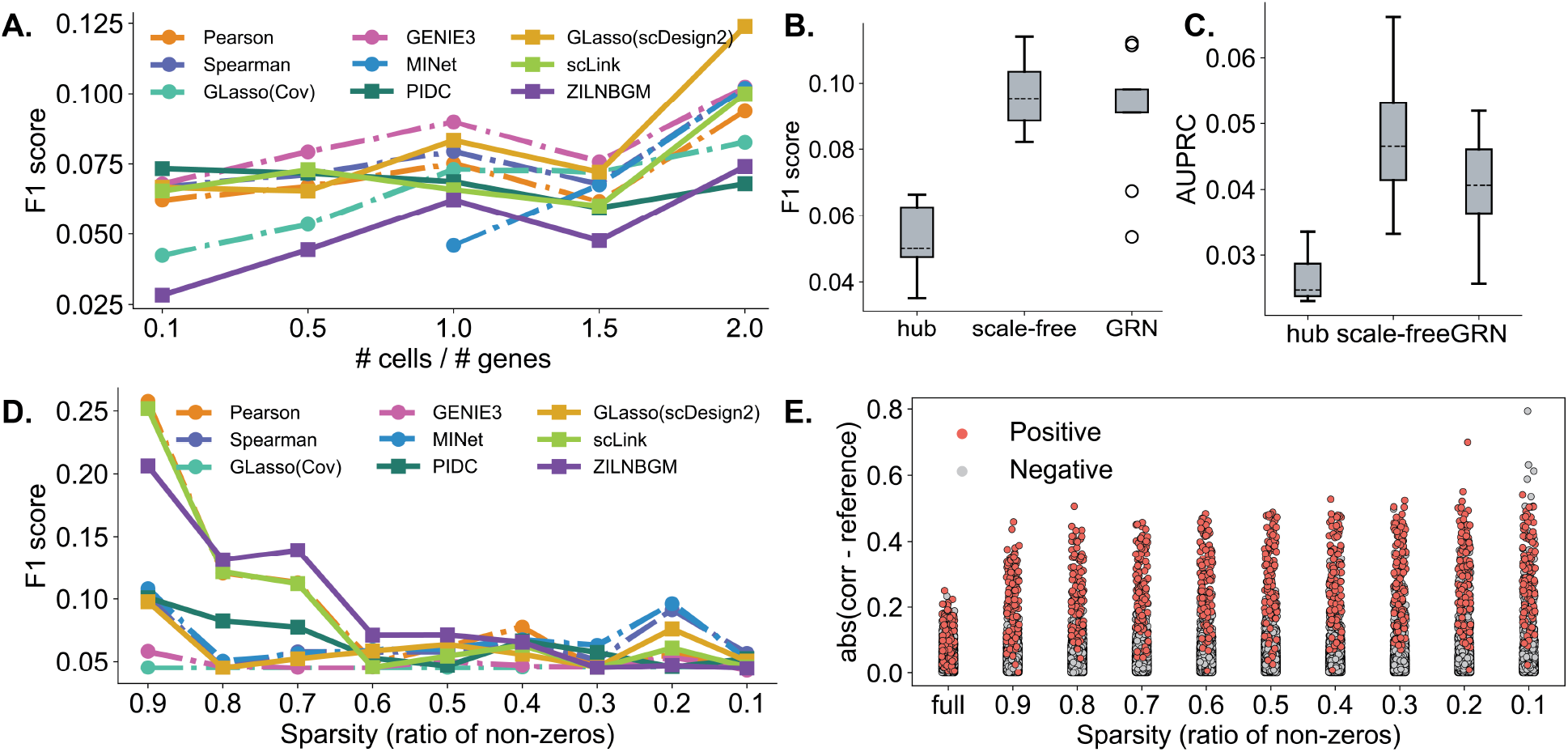
Methods performances on simulated data with different settings. (**A**) F1-scores of `NORTA` simulations with 100 genes and {10, 50, 100, 150, 200} cells. (**B-C**) F1-scores and AUPRC values of methods on `NORTA` simulations with three different graph structures. (**D**) F1-scores of `NORTA` simulations with different sparsity levels. (**E**) The absolute difference between simulation correlations and the true network at different sparsity levels.

Last, we simulate `NORTA` data with different sparsity levels using negative binomial gene marginal distributions and adding zero counts following Splatter simulation (Zappia et al., 2017). We test methods on data with 100 genes and {10%, 20%, …, 90%} sparsity. Fig. 9(D) implies that the estimation worsens as the data becomes more and more sparse. To understand how sparsity affects co-expression estimation, we compute the Pearson correlation coefficients of data at different sparsities and study how the correlation changes. Fig. 9(E) computes the absolute difference between the simulation correlation and the true network. It shows that when data gets sparser, the simulation correlation becomes more different from the true network due to increased heterogeneity in the data, resulting in difficulties in co-expression evaluation. The increase in difference involving both positive (i.e., co-expression in the true network) and negative (i.e., non-co-expression in the true network) edges in the true network is perhaps the reason for the high FDR. Our results indicate that the sparsity in data is the major bottleneck of estimation and methods that can address sparsity should be investigated for future co-expression estimation method development.

### 3.6 Data Imputation or Pre-Processing steps introduce More False Positives in co-expression estimation

In scRNA-seq analysis, many pre-processing and imputation approaches have been proposed to address the analytical difficulties arising from the sparsity of the data. Here, we apply a few commonly used pre-processing and imputation approaches to test whether they can improve the co-expression estimation.

First, normalization is a critical pre-processing step in scRNA-seq analysis to adjust unwanted biases and obtain information of interest (Grün and van Oudenaarden, 2015; Vallejos et al., 2017; Chen et al., 2019). We utilize three commonly-used and novel normalizing schemes: `LogNorm` (Satija et al., 2015; Zheng et al., 2017; Stuart et al., 2019; Butler et al., 2018), `sctransform` (Hafemeister and Satija, 2019; Choudhary and Satija, 2022), and `PsiNorm` (Borella et al., 2022). Details about these normalization methods are provided in **Appendix: More on Methods and Results**. After comparing method estimations from the raw expression matrix and normalized matrix, we find the normalized data does not improve and even worsens the method performance (Fig. 10(A-B)) on both types of simulations. Appendix Table A2 indicates that `LogNorm` and `PsiNorm` cannot reduce the sparsity in data such that challenges of estimation remain. For `sctransform`, it is designed to remove variations caused by RNA sequencing depths. But there are other types of technical variations (Haque et al., 2017), such as cell cycle (Buettner et al., 2015) and transcriptional bursting (Suter et al., 2011). So the reason `sctransform` does not improve the estimation is potentially due to its lack of ability to address multiple types of variations.

**Figure 10:**
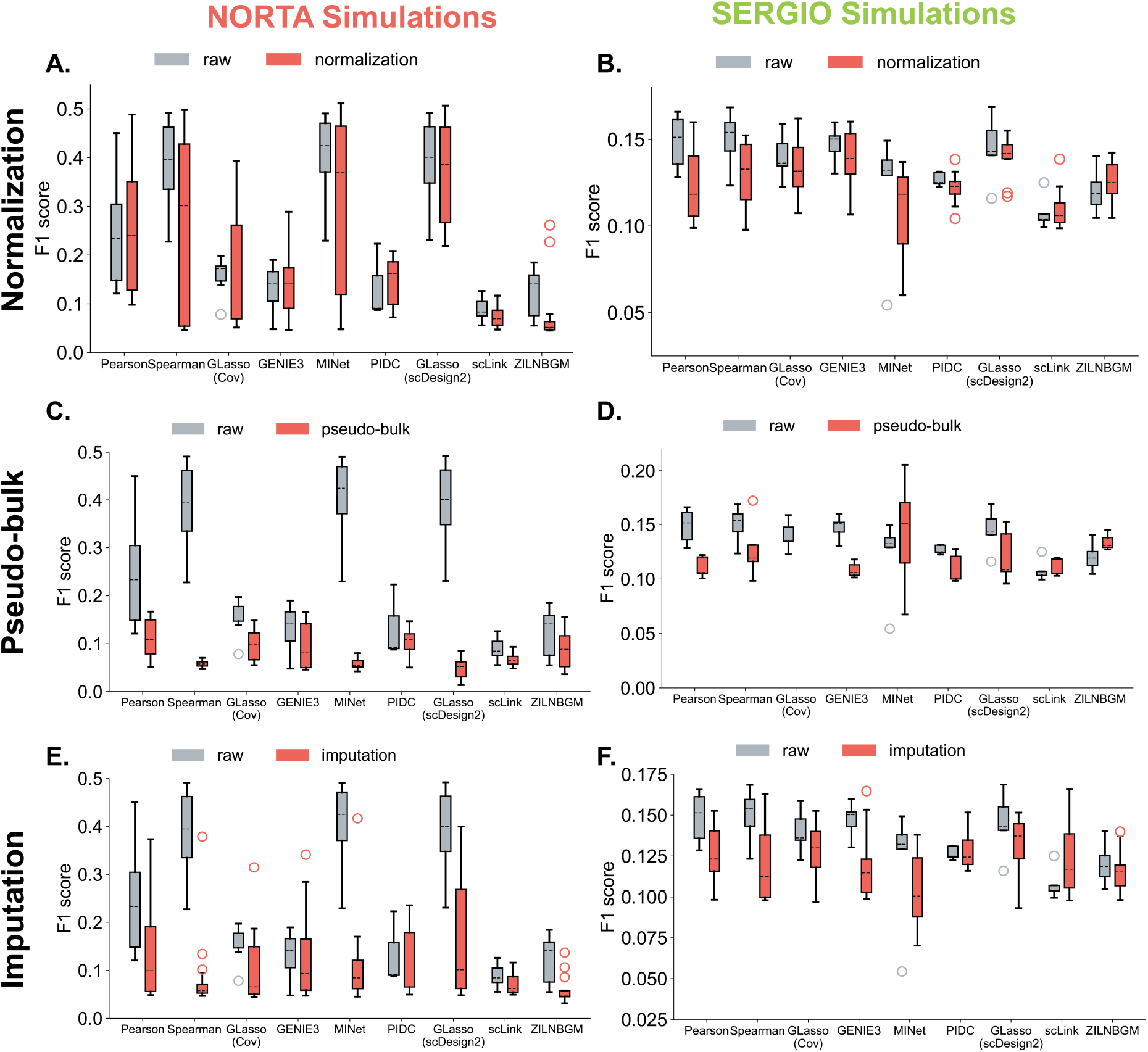
Comparison of methods performances on `NORTA` simulations and corresponding processed data from different approaches. (**A-B**) F1-scores for raw and normalized simulated data from `LogNorm`, `PsiNorm`, and `sctransform`. (**C-D**) F1-scores for raw simulations and pseudo-bulks. (**E-F**) F1-scores for raw and imputed simulations from `MAGIC`, `SAVER`, `DCA`, and `ALRA`.

One other way to reduce variations in scRNA-seq data is calculating the pseudo-bulk signal (Squair et al., 2021), which is obtained from averaging cell expressions for cells in the same cluster. For each simulation dataset, we apply UMAP to find the low-dimensional representation and then cluster cells with Leiden clustering. The pseudo-bulk expression is obtained by averaging cell expressions in each cluster. Fig. 10(C-D) show that the pseudo-bulk cannot improve method predictions and might break down the method, e.g., for `MINet`. We hypothesize that clustering may induce biases, and constructing the pseudo-bulk results in statistical information loss, which affects the co-expression estimation.

To solve the problem with sparsity in scRNA-seq data, imputation (Hou et al., 2020) is a common solution. In scRNA-seq data, observed zeros can be either biological zeros denoting non-expression of genes or a technical limitation in quantifying the small number of molecules. To address the challenges induced by the sparsity, some recent works have developed several imputation methods to impute observed zeros. They were shown to improve downstream single-cell data analysis (Hou et al., 2020), such as differential gene expression analysis and cell clustering. Here, we test whether the imputation can improve gene co-expression estimation. We apply all methods on simulation data imputed with four commonly used state-of-the-art algorithms, `MAGIC`(Van Dijk et al., 2018), `SAVER`(Huang et al., 2018), `DCA` (Eraslan et al., 2019), and `ALRA` (Linderman et al., 2022). Appendix Table A2 shows the imputation has reduced data sparsity. But results indicate that imputed data worsen the method predictions, producing higher FDR scores (Fig. 10(E-F) and Appendix Fig. A14). The imputation step has been reported to induce artificial correlations in data (Andrews and Hemberg, 2018; Hou et al., 2020) and thus might explain the observed reduced estimation performance.

In addition, simply removing lowly-expressed cells seems useful. So for each `NORTA` simulated dataset, we remove cells with library size smaller than *t*% × *K, t* ∈ {1%, 5%, 10%, 15%, 20%}, in which *K* is the maximal library cell of all cells. Appendix Fig. A18 shows that removing lowly-expressed cells doesn’t improve performance either. This is reasonable because Appendix Fig. A1 shows that all cells are sparse, so simply removing cells will not reduce data sparsity.

Overall, we have found that these commonly used pre-processing approaches do not improve the gene co-expression estimation and might further introduce more false positives, resulting in estimations of lower quality.

## 4 Discussion and Conclusion

In this study, we test several gene co-expression estimators on scRNA-seq data. Experiment results show that existing methods tend to produce estimations with high FDR. The choice of realistic underlying assumptions and testing on simulations that represent properties of experimental data are keys to developing robust estimation methods. We find that methods like `GLasso` and `GENIE3` that are proposed for bulk data do not perform well for scRNA-seq data because scRNA-seq data distribution is significantly different. We also show that previously used simulations, `ZI-Gaussian` and `ZI-Poisson`, have much fewer zero counts and different distribution shapes compared to real-world experimental data. Therefore, we propose the use of two types of new simulated datasets, `NORTA` and `SERGIO`, that exhibit characteristics of experimental data. The low performance of existing methods on these simulations needs to be considered when applying them to real-world single-cell datasets. Especially a co-expression estimation with a low FDR is required in the field.

We also test different pre-processing steps of scRNA-seq datasets to see whether they can improve the estimation. We find that the widely-used normalization step can reduce data variations, however, the high-level of sparsity in the data continues to make co-expression estimation challenging. Next, we use pseudo-bulk to reduce data sparsity by averaging cell expressions through clustering. But it obtains a worse estimation compared to raw data in our experiments. We hypothesize that the clustering may induce biases in the data. Furthermore, constructing the pseudo-bulk reduces the number of samples and results in statistical information loss, which negatively affects the analysis. Finally, we show that scRNA-seq imputation methods also worsen the methods’ performance by potentially inducing artificial correlations in data. Overall, the pre-processing steps commonly used on scRNA-data do not help with co-expression estimation due to various limitations and biases related to the complexity of single-cell datasets.

While our study shows that none of the existing methods provide correct estimations, we believe a few methods can still be used practically as first steps for downstream improvement or analysis. Based on our experiments, we recommend using `Spearman`, `MINet`, or `GLasso` with `scDesign2` transformation to measure gene co-expression since they have better performance in our experiments.

Unsurprisingly, we find that the main challenge of scRNA-seq data is its high sparsity. This makes robust analysis of the data a difficult task. We suggest that, in future works, the improvement on gene co-expression estimation should be made upon model robustness against sparsity and FDR control. Another potential solution might be adding more data from other modalities to help with the estimation. Due to the high sparsity and variations, the scRNA-seq data itself may not provide enough information for promising analysis, so integrating other data, for example scATAC-seq data, perhaps is important for more comprehensive and confident analyses.

In summary, our tested simulations will be helpful in gene co-expression analysis, especially for benchmarking and validating methods. We think our study provides valuable insights for scRNA-seq data analyses, and our benchmark setup can be a resource for the community as we work towards developing more accurate gene co-expression estimators.

## Code Availability

All the codes and data used in this study are available at https://github.com/rsinghlab/scRNAseq_Coexpression_Benchmark.

### Acknowledgements

We thank Pinar Demetci and Tassallah Amina Abdullahi for their useful comments on this study.

## A1 More on Methods and Results

### A1.1 Properties Comparison between Simulated and Experimental Data

In Sec. 3.1 and 3.2, we compare several basic statistics between simulated and experimental scRNA-seq data. These statistics include the percent of zeros per cell, percent of zeros per gene, distribution shape, and cell clusters.

In comparisons, we use box plots for the percent of zeros per cell/gene. The similarity between the two datasets is evaluated via the Kolmogorov–Smirnov test (KS test), where the null hypothesis is that the sparsity of the two datasets is identical. To compare distribution shapes, since different dataset has different scales of values, we normalize the data into range [0.0, 1.0] with min-max scaling

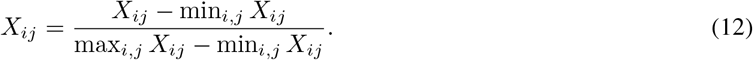

We compare distribution shapes qualitatively through frequency histograms and quantitatively through integrated absolute error (IAE) as

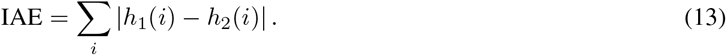

Here *h*_1_ and *h*_2_ are 20-bins histograms of two datasets, and *h*_1_(*i*) is the data frequency at the *i*-th bin.

We use clustering to validate that simulations have the multi-cell-type structure as experimental data. Concretely, given an expression matrix **X** ∈ ℝ*n*×*p*, we apply Principal Component Analysis (PCA) to find the top 50 principal components as **P** = PCA_50_(**X**) ∈ ℝ*n*×50. Then we utilize the Leiden clustering (Traag et al., 2019) to split cells into clusters from **P**. We visualize the clustering results in the two-dimensional Uniform Manifold Approximation and Projection (UMAP) (McInnes et al., 2018) space **U** = UMAP_2_(**P**) ∈ ℝ*n*×2. Moreover, we use the silhouette score (Rousseeuw, 1987) to evaluate the clustering performance. Specifically, let 𝒞_*k*_ denote a set of cells in the *k*-th cluster, the silhouette score is computed through

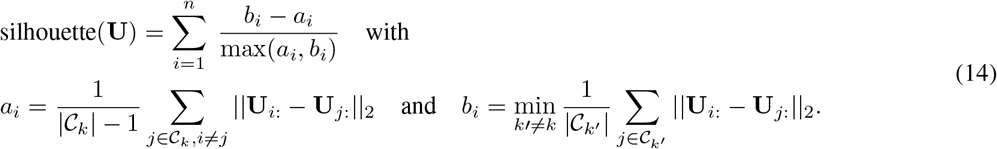

Here, *a*_*i*_ measures the average ℓ_2_ distance between a cell *i* and every cell in the same cluster. *b*_*i*_ measures the average ℓ_2_ distance between cell *i* and every cell in the nearest cluster. Appendix Fig. A2, A3, and A4 provide clustering results along with silhouette scores for 100 HVGs experimental data and `NORTA`/`SERGIO` simulations. We use `scanpy` (Wolf et al., 2018) for PCA, UMAP, and Leiden clustering. We use `scikit-learn` (Pedregosa et al., 2011) to compute the silhouette score.

To validate that our new `NORTA` simulation can successfully keep the given gene co-expression structure, we compare the correlation of data before and after tuned by a given network. Let **W**\* denote the true covariance matrix used in `NORTA` simulation; Corr_exp_ denote Pearson correlation coefficient matrix of experimental data; Corr_sim_ denote correlation matrix of corresponding simulated data. The Corr_sim_ should be closer to **W**\* since we explicitly impose the network structure in `NORTA`. So we utilize the correlation matrix distance (CMD) (Herdin et al., 2005) to measure the similarity between two correlation matrices as

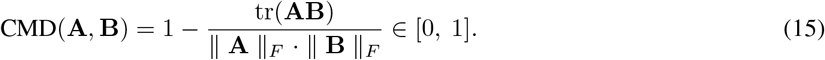

CMD(**A, B**) = 0 denotes two correlation matrices are equal up to a scaling factor and CMD(**A, B**) = 1 if they differ in max extents. Therefore, CMD(Corr_sim_, **W**\*) *<* CMD(Corr_exp_, **W**\*) indicates that the simulation successfully impose the gene network structure on data. Apart from CMD, we also utilize mean squared error (MSE) as

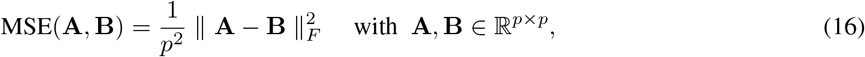

in which ‖ · ‖_*F*_ is Frobenius norm and a smaller MSE value indicates two matrices are more similar.

### A1.2 Gene Co-Expression Network Estimators

- `Pearson`/`Spearman`: The correlation coefficient is the most intuitive measure of gene co-expression (Langfelder and Horvath, 2008). In this study, we test `Pearson` and `Spearman` correlation. Specifically, for every pair of genes, we compute the correlation coefficients and constructed a correlation matrix as

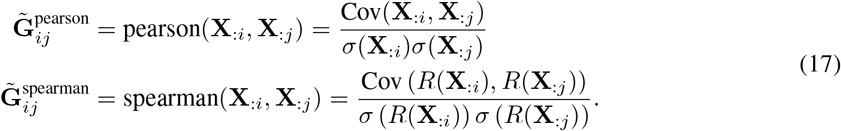

Here Cov is the covariance, *Σ* is the standard deviation, and *R*(**a**) : ℝ*n* ↦ ℝ*n* are ranks of variable **a**. Then we filter the correlation matrix 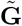 with a user-defined threshold *δ* for the final estimation as 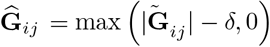

- `MINet`: Mutual information (MI), as a generalized correlation measure, has also been used to estimate gene co-expression networks. The MI is estimated with Spearman entropy estimator, and the network is estimated through MRNET (Meyer et al., 2007) algorithm. We also use *δ* to threshold the mutual information matrix. This method is referred to `MINet` in this study. We use R package `minet` (Meyer et al., 2008) in our implementation.
- `GLasso(Cov)`: (Friedman et al., 2008) proposed graphical lasso (GLasso) to estimate a sparse inverse covariance matrix from Gaussian data. Specifically, GLasso assumes *p* genes follow a multivariate Gaussian distribution 𝒩 (**0**, Σ) and the inverse covariance **G** := Σ^−1^ encodes the gene co-expression network. We generally solve GLasso with a maximum likelihood estimation (MLE)

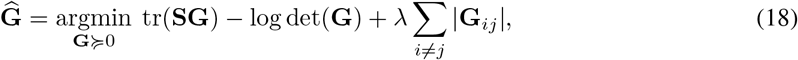

in which **S** is the sample covariance matrix and *λ* is a hyper-parameter controls sparsity. We name it `GLasso(Cov)` because it uses GLasso and is based on the covariance matrix. We use the implementation of R package `QUIC`(Hsieh et al., 2011).

- `GENIE3`: (Huynh-Thu et al., 2010) developed `GENIE3` to estimate gene regulatory network from microarray data. The essential idea of `GENIE3` is decomposing the task of estimating a network of *p* genes into *p* subproblems, each aiming to estimate one gene’s interactions based on regression. Each subproblem is solved with a tree-based ensemble method. We use R package `GENIE3` in our implementation.
- `PIDC`: (Chan et al., 2017) developed `PIDC` to estimate gene regulatory network from single-cell data. It utilizes partial information decomposition to estimate relationships between genes. We use R package `PIDC` in our implementation.
- `scLink`: (Li and Li, 2021) proposed a gene co-expression estimator, `scLink`, for scRNA-seq data. It assumes log-transformed scRNA-seq data follow the mixture of Gamma-Normal distribution. Upon this, `scLink` estimate robust correlation coefficients to ease the influence of excess zeros in data. The robust correlation matrix is later fed into GLasso to give a final gene co-expression network estimation. we follow the `scLink` R implementation in the study.
- `GLasso(scDesign2)`: (Sun et al., 2021) originally proposed `scDesign2` for simulating high-fidelity scRNA-seq data and it can also be used on gene co-expression estimation. It assumes the expression of each gene follows zero-inflated negative binomial (ZINB) distribution. It first converts each gene’s expression to Gaussian-like samples with a Gaussian copula. Then it simulates data with the approximated covariance matrix of transformed data. In our study, we take this covariance matrix of transformed data from `scDesign2` and use GLasso to estimate gene co-expression.
- `ZILNBGM`: (Choi et al., 2017; Park et al., 2021) estimated gene co-expression network with the local graphical model. For each node (i.e., gene), it locally infers each node’s neighbors with zero-inflated negative binomial regression and integrates neighbors of all nodes into a gene network. It has a hyper-parameter *λ* to control the sparsity. This approach is named zero-inflated local negative binomial graphical model (`ZILNBGM`).

### A1.3 scRNA-seq Simulation with Different Settings

- **Different number of cells:** We use `NORTA` to generate five datasets of {10, 50, 100, 150, 200} cells corresponding to 100 HVGs subsets of eight experimental datasets. We use the same hub-based co-expression network in these simulations. The number of cells ranges from 0.1 × to 2× of the number of genes, providing a sequence of tests from low-dimensional (where the number of cells *n* are larger than the number of genes *p*) to high-dimensional settings (where *n < p*). Totally, we have 40 simulated datasets in this case.
- **Different network structures:** We use `NORTA` to generate data based on three different co-expression graph structures: hub-based, scale-free, and GRN. We use `pcorSimulator` R package to generate hub-based and scale-free networks. The GRN network is taken from example data in (Dibaeinia and Sinha, 2020). The gene distributions are fitted from 100 HVGs subsets of eight experimental datasets, so we have 24 simulated datasets in this case.
- **Different sparsity level:** We simulate `NORTA` data of different sparsity. We use the hub-based network and a negative binomial distribution with dispersion= 0.5 and probability of success= 0.005 to generate expression matrix **X** of 100 genes and 500 cells. Then we add zero counts on **X** following Splatter simulation (Zappia et al., 2017), to obtain data of {10%, 20%,, 90%} sparsity. Concretely, let **Y** = [log(**X**_*ij*_ + 1)]*i,j*, we add zeros through

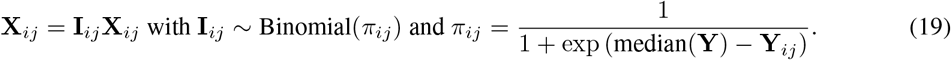

### A1.4 scRNA-seq Simulation with Different Pre-processing Approaches

In Sec. 3.6, we test multiple processing approaches to validate whether they can improve the co-expression estimation.

- **Normalization**: We test three normalizing techniques:
- **LogNorm**: Specifically, given a count matrix **X** ∈ ℝ*n*×*p* of *n* cells and *p* genes, we first normalize each cell with its library size as

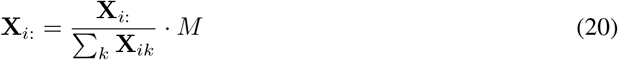

where *M* is a scale factor and we set it as *M* = 10^4^ in our experiments. After this, we log-transform data with

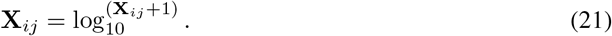

- **sctransform**: (Hafemeister and Satija, 2019; Choudhary and Satija, 2022) proposed a novel scRNA-seq data normalization method called `sctransform`. It is designed to stabilize cell sequence depth variations, and it models gene expression with the generalized linear model (GLM). Concretely, `sctransform` assumes the expression of gene *j* at cell *i* follows

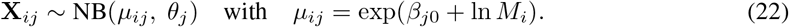

Here 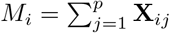 is the library size of cell *i*. `sctransform` fits the NB distribution for each gene through the regularized negative binomial regression. Then `sctransform` normalizes data by converting it to the Pearson residual via

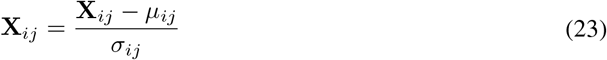

with *μ*_*ij*_ = exp(*β*_*j*0_ + ln *M*_*i*_) and 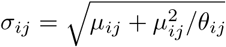. Note that `sctransform` normalized data have negative values. So `ZILGM` and `GLasso(scDesign2)` are infeasible for `sctransform` outputs since they require data to be non-negative. We use R package `sctransform` in our experiments.

- **PsiNorm**: (Borella et al., 2022) proposed a scalable normalization method `PsiNorm` for scRNA-seq data, based on power-law Pareto distribution. Specifically, for a cell *i*, `PsiNorm` normalizes its expression through

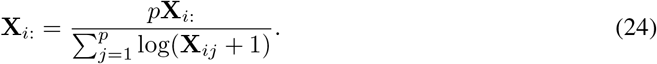

We use R package `PsiNorm` in our experiments.

- **Pseudo-bulk**: Pseudo-bulk data are obtained in two steps. First, we cluster cells with Leiden clustering on their UMAP representations. In UMAP, The neighbor graph is computed with the number of neighbors as ten on the first 50 principal components. Then for each cluster, we average cell expressions to get the final pseudo-bulk expression matrix. We use `scanpy` package (Wolf et al., 2018) to obtain pseudo-bulk data in implementation. Appendix Table A2 shows the pseudo-bulk data sparsity.
- **Data Imputation**: We also test how scRNA-seq imputation algorithms affect co-expression estimation since they can reduce the data sparsity. We utilize four imputation algorithms, `MAGIC` (Van Dijk et al., 2018), `SAVER` (Huang et al., 2018), `DCA` (Eraslan et al., 2019), and `ALRA` (Linderman et al., 2022) to impute simulation datasets. `MAGIC` imputes scRNA-seq data through diffusion on the cell similarity network. Specifically, `MAGIC` first compute distances between every pair of cells and normalize them into cell affinities. The imputation is completed by performing diffusion on cell affinities. `SAVER` first learns the posterior distribution of each gene from regression methods and imputes zeros from learned distributions. `DCA` takes overdispersion and sparsity into consideration and utilizes a variational autoencoder to denoise the data. These three methods are commonly used in scRNA-seq data analysis. Moreover, the recently proposed `ALRA` impute data while preserving biologically non-expressed values based on low-rank matrix approximation. Our study runs each imputation algorithm on simulated datasets with default hyper-parameter settings. Appendix Table A2 shows the data sparsity after imputation.

### A1.5 Gene Co-expression Estimation On Simulations with More Genes

We also test all methods on `NORTA` simulations with 500 genes and `SERGIO` simulations with 400 genes. We use `NORTA` to generate data corresponding to 500 HVGs of eight experimental data with hub-based co-expression networks. For `SERGIO` simulation, we use the example data of 400 genes in (Dibaeinia and Sinha, 2020). Appendix Fig. A7 and A8 show that, compared with simulations with 100 genes (Fig. 7), methods perform worse on data with more genes. This indicates that more genes worsen the estimation due to increased problem complexity.

### A1.6 Gene Co-expression Estimation On Experimental Data of Different Cell Types

To remove the mutual effects of multiple cell types, We follow (Li and Li, 2021) and evaluate all methods on gene expressions of individual cell types. Concretely, we use Tabula Muris database (Consortium et al., 2018), which is derived from Smart-seq2 RNA-seq libraries (Picelli et al., 2014) and contains 53,760 cells from 20 tissues of 8 mice. We use expressions of 500 HVGs from three cell types: T cells (793 cells and sparsity=7.37%), pancreatic beta cells (449 cells and sparsity=22.59%), and skeletal muscle satellite stem cells (439 cells and sparsity=7.24%). We still compare mouse TF co-expression networks with method predictions. Appendix Fig. A19 shows that all methods have average F1-scores*<*0.4. `MINet` performs the best on recovering known TF co-expression with PPV ≈ 0.41. Other methods have PPV*<* 0.3.

## A2 Supplementary Figures

**Figure A1:**
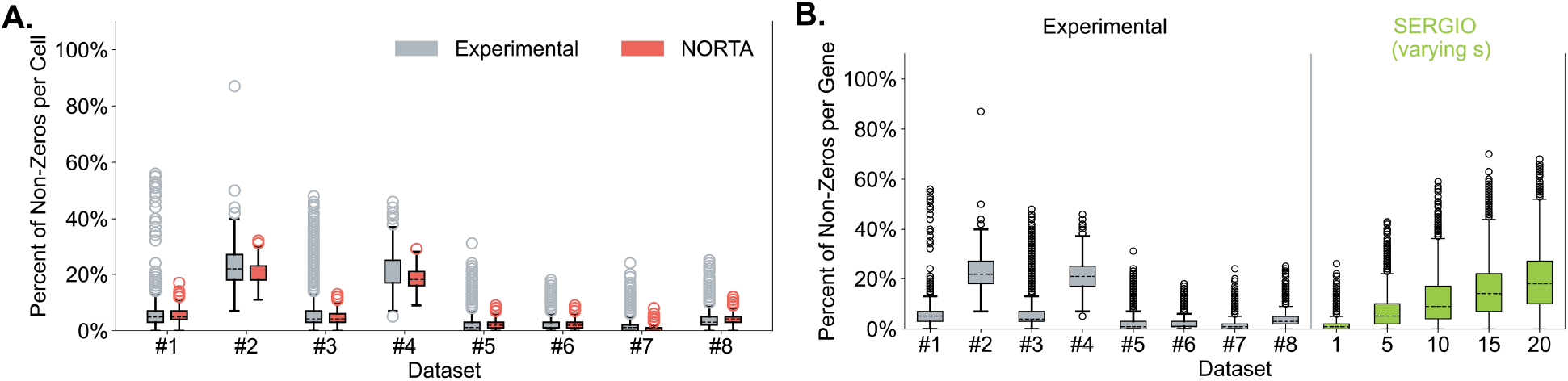
Compare experimental data with `NORTA` and `SERGIO` simulations concerning cell sparsity. (**A**) Cell sparsity of experimental data and corresponding `NORTA` simulation. (**B**) Cell sparsity of experimental data and `SERGIO` simulations varying *s* ={1, 5, 10, 15, 20}.

**Table A1:**
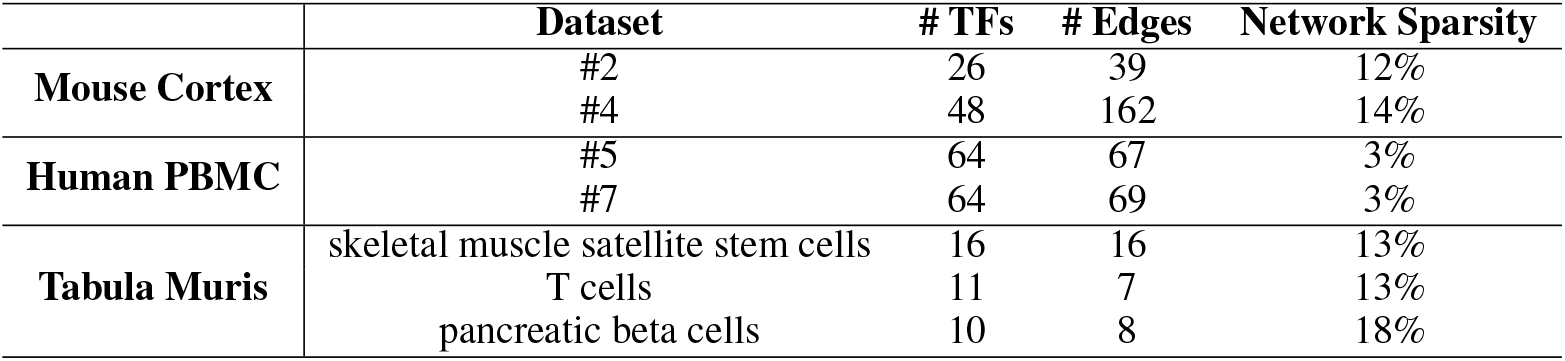
Basic stats of TF co-expression networks.

**Figure A2:**
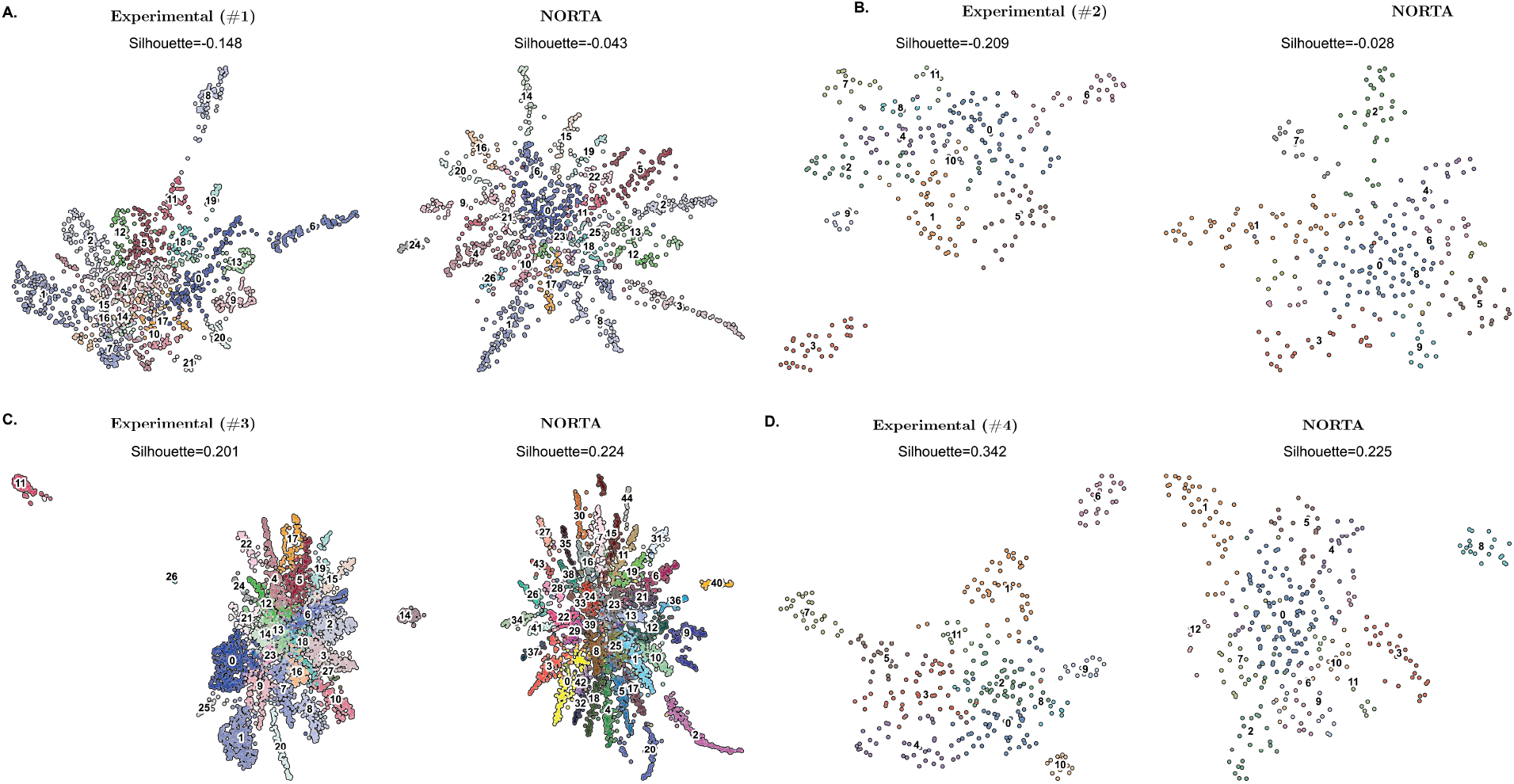
Compare cell clustering on mouse cortex experimental data (#1 to #4) and corresponding `NORTA` simulations.

**Figure A3:**
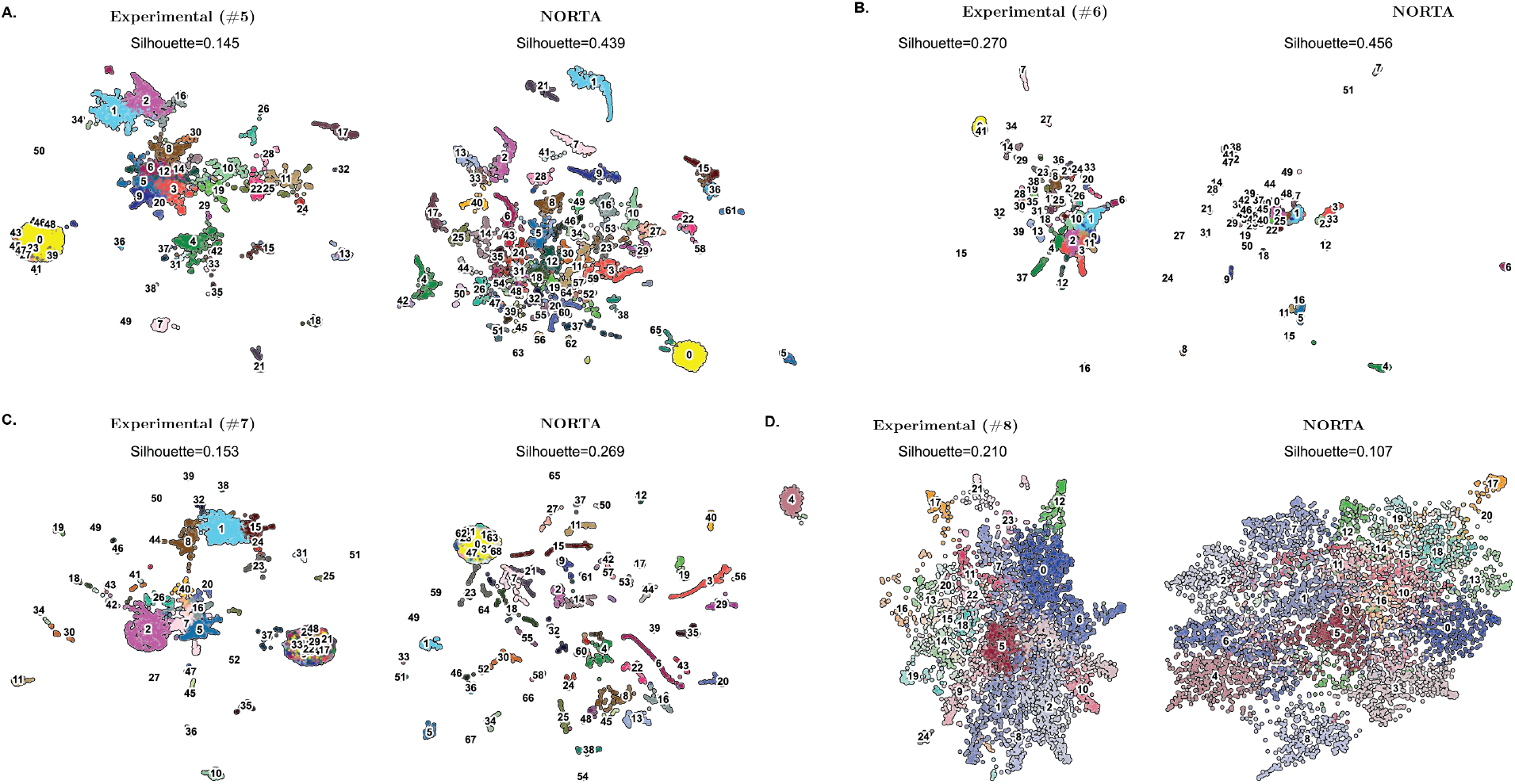
Compare cell clustering on PBMC experimental data (#5 to #8) and corresponding `NORTA` simulations.

**Figure A4:**
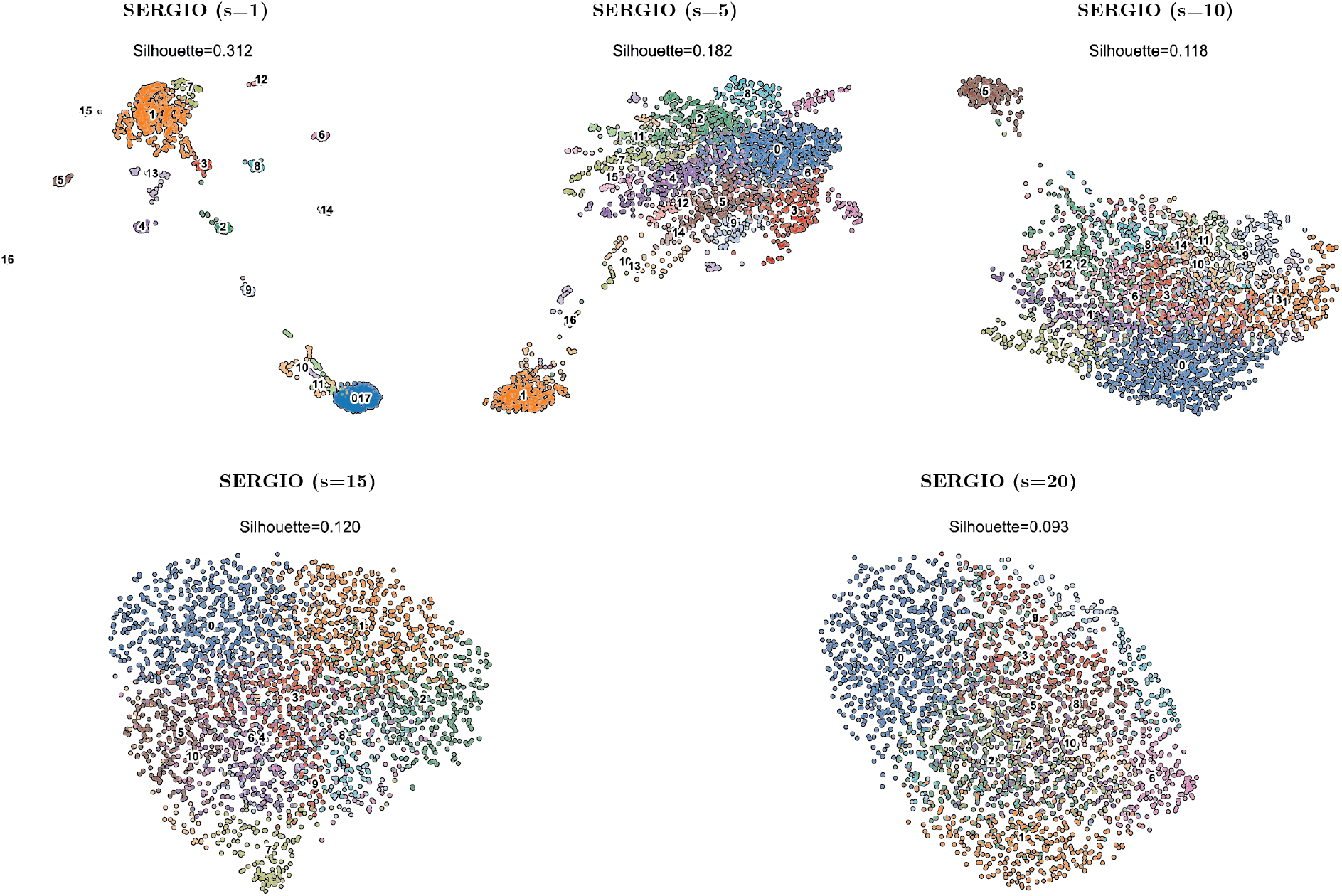
Cell clustering of `SERGIO` simulations with s={1, 5, 10, 15, 20}.

**Figure A5:**
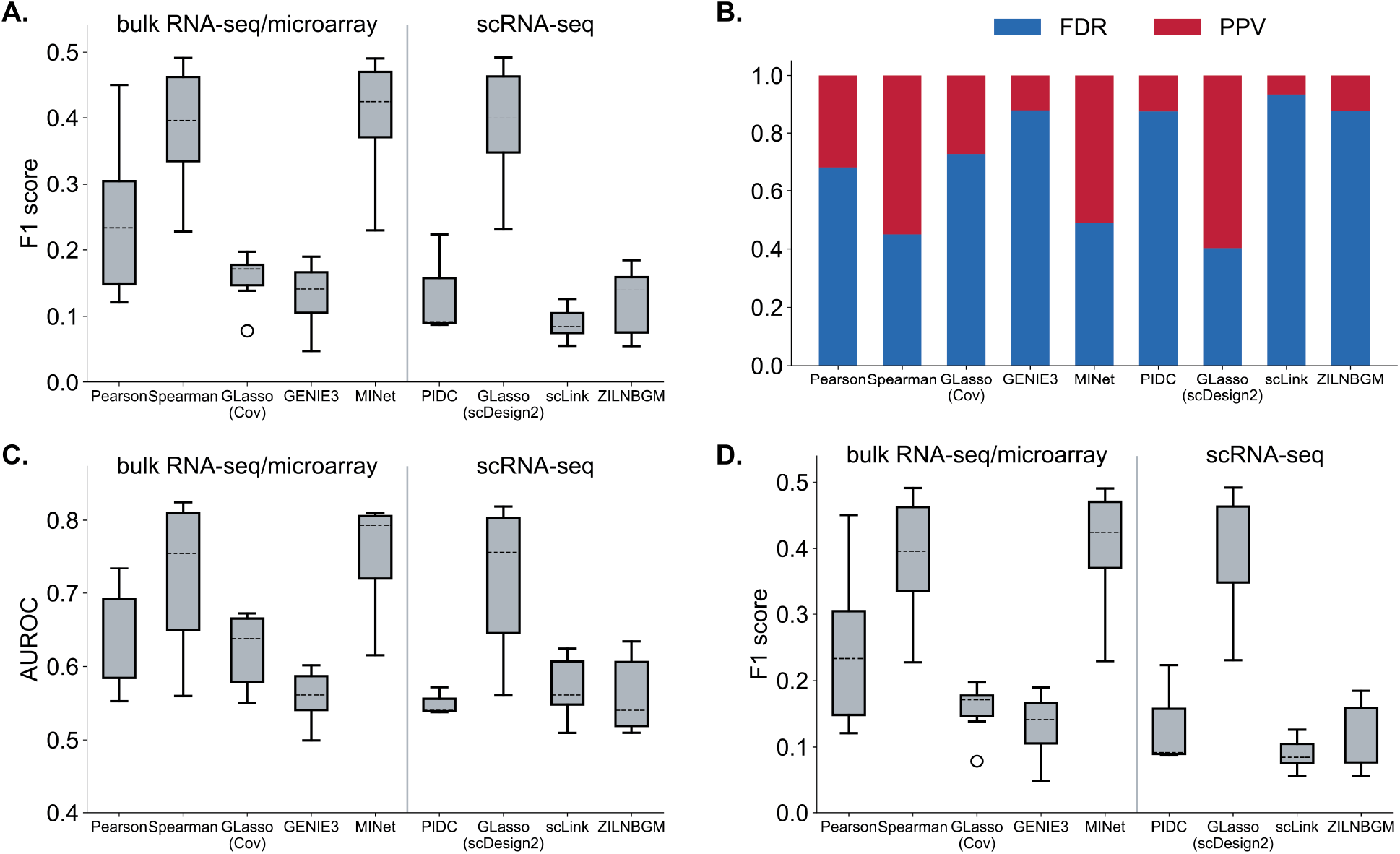
Method performances on eight `NORTA` simulations with 100 genes, including (**A**) boxplot for F1-score, (**B**) FDR and PPV, (**C**) boxplot for AUROC, and (**D**) boxplot for AUPRC.

**Figure A6:**
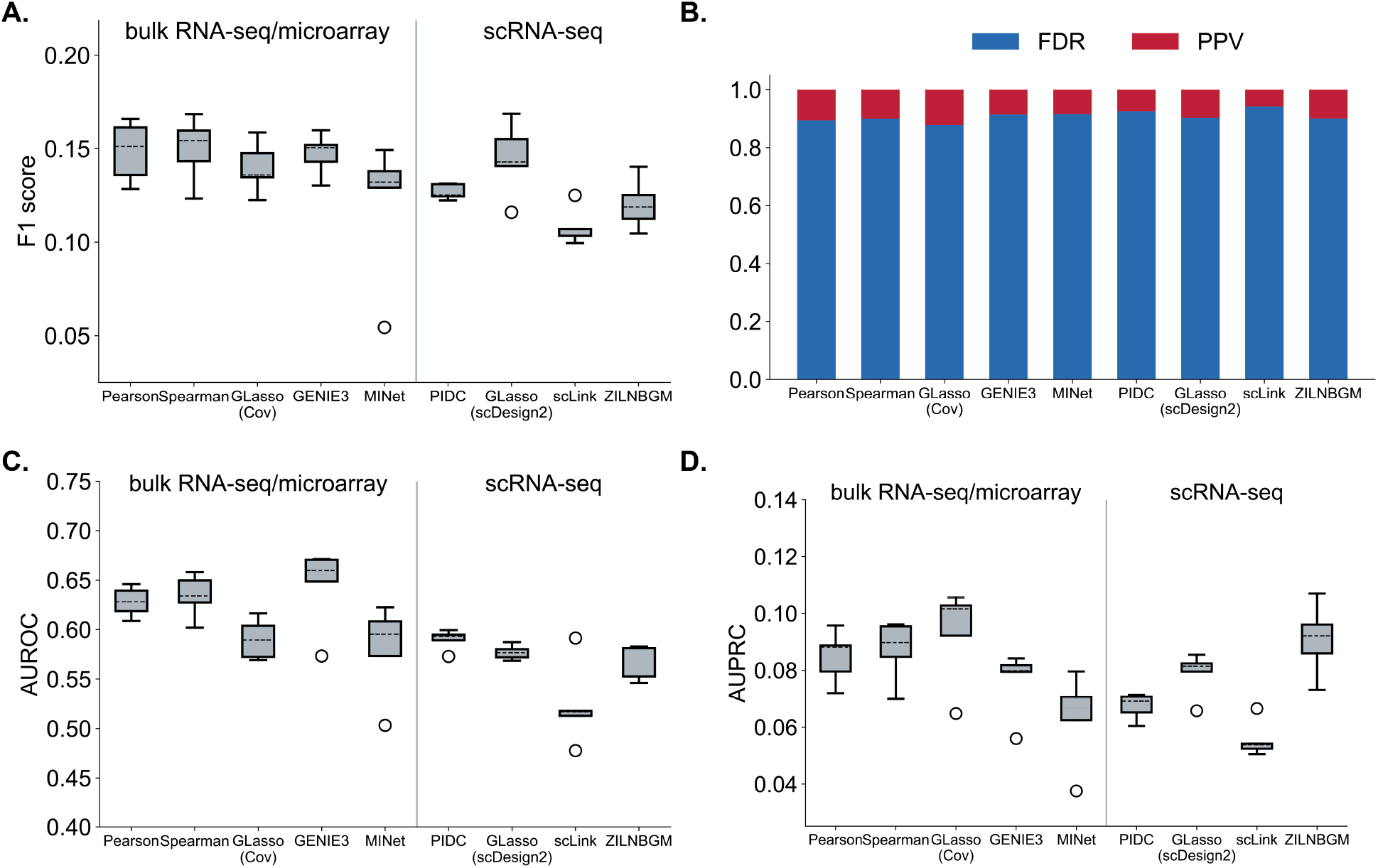
Method performances on five `SERGIO` simulations with 100 genes, including (**A**) boxplot for F1-score, (**B**) FDR and PPV, (**C**) boxplot for AUROC, and (**D**) boxplot for AUPRC.

**Table A2:**
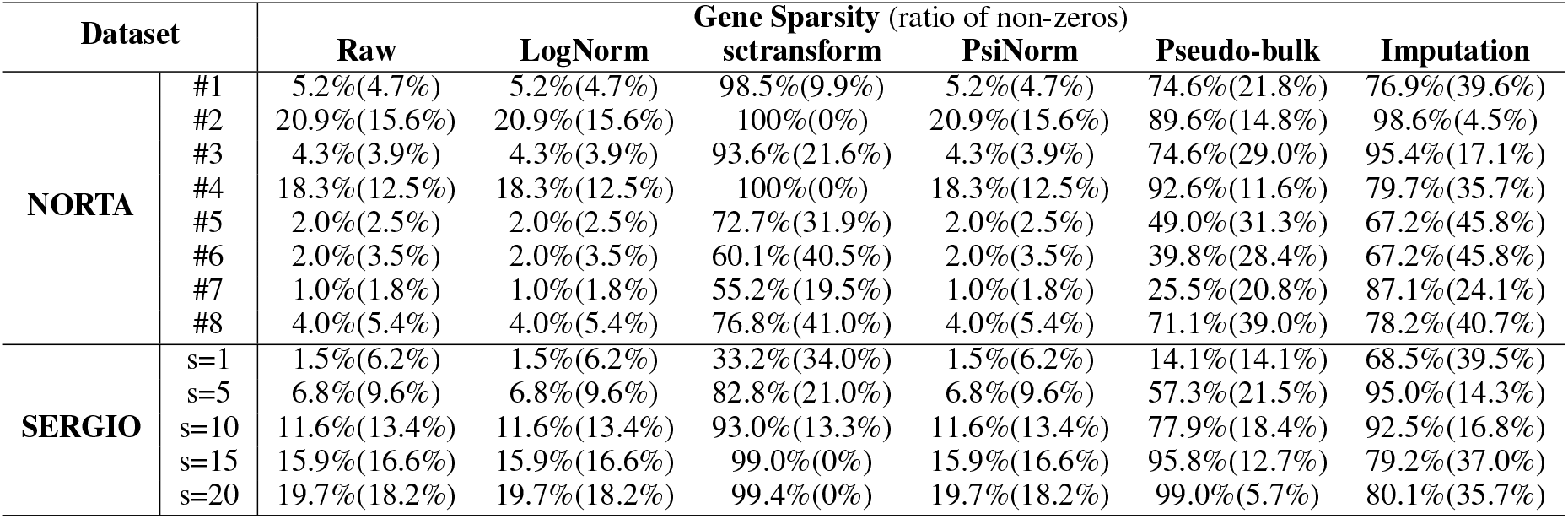
Average(std.) gene sparsity (ratio of non-zeros) of 100HVG simulated datasets of raw data and different pre-processed data.

**Figure A7:**
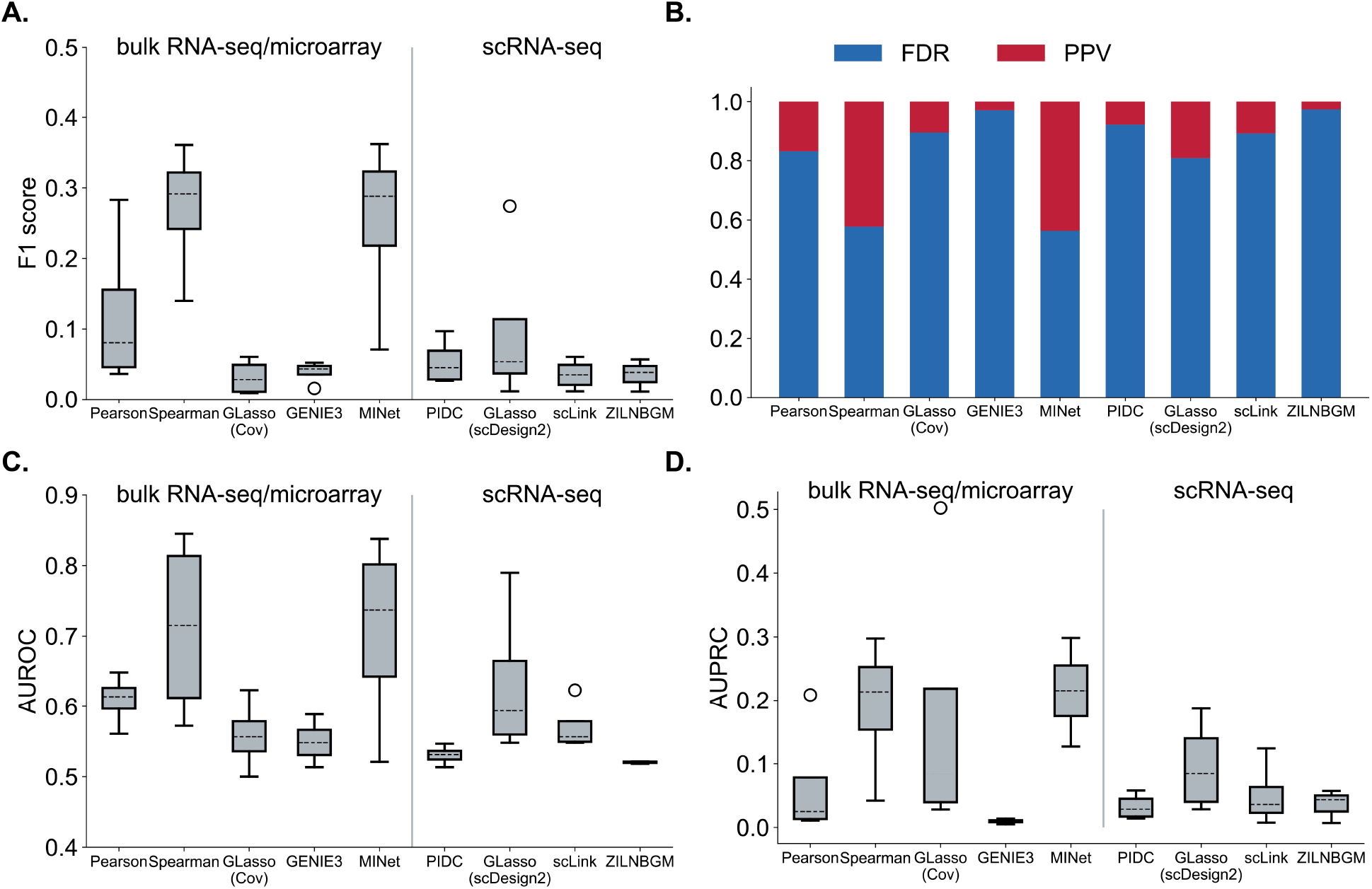
Method performances on eight `NORTA` simulations with 500 genes, including (**A**) boxplot for F1-score, (**B**) FDR and PPV, (**C**) boxplot for AUROC, and (**D**) boxplot for AUPRC.

**Figure A8:**
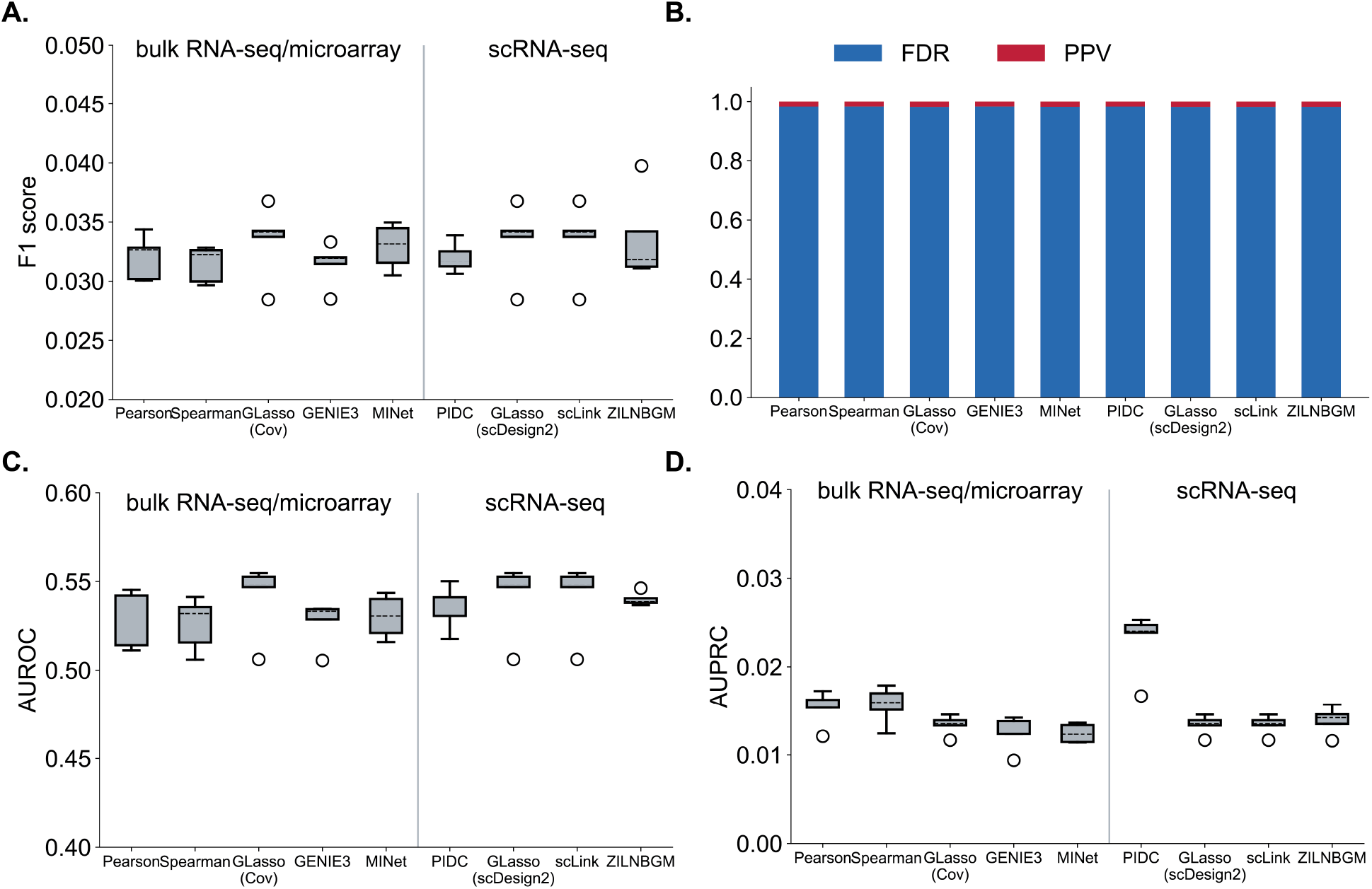
Method performances on five `SERGIO` simulations with 400 genes, including (**A**) boxplot for F1-score, (**B**) FDR and PPV, (**C**) boxplot for AUROC, and (**D**) boxplot for AUPRC.

**Figure A9:**
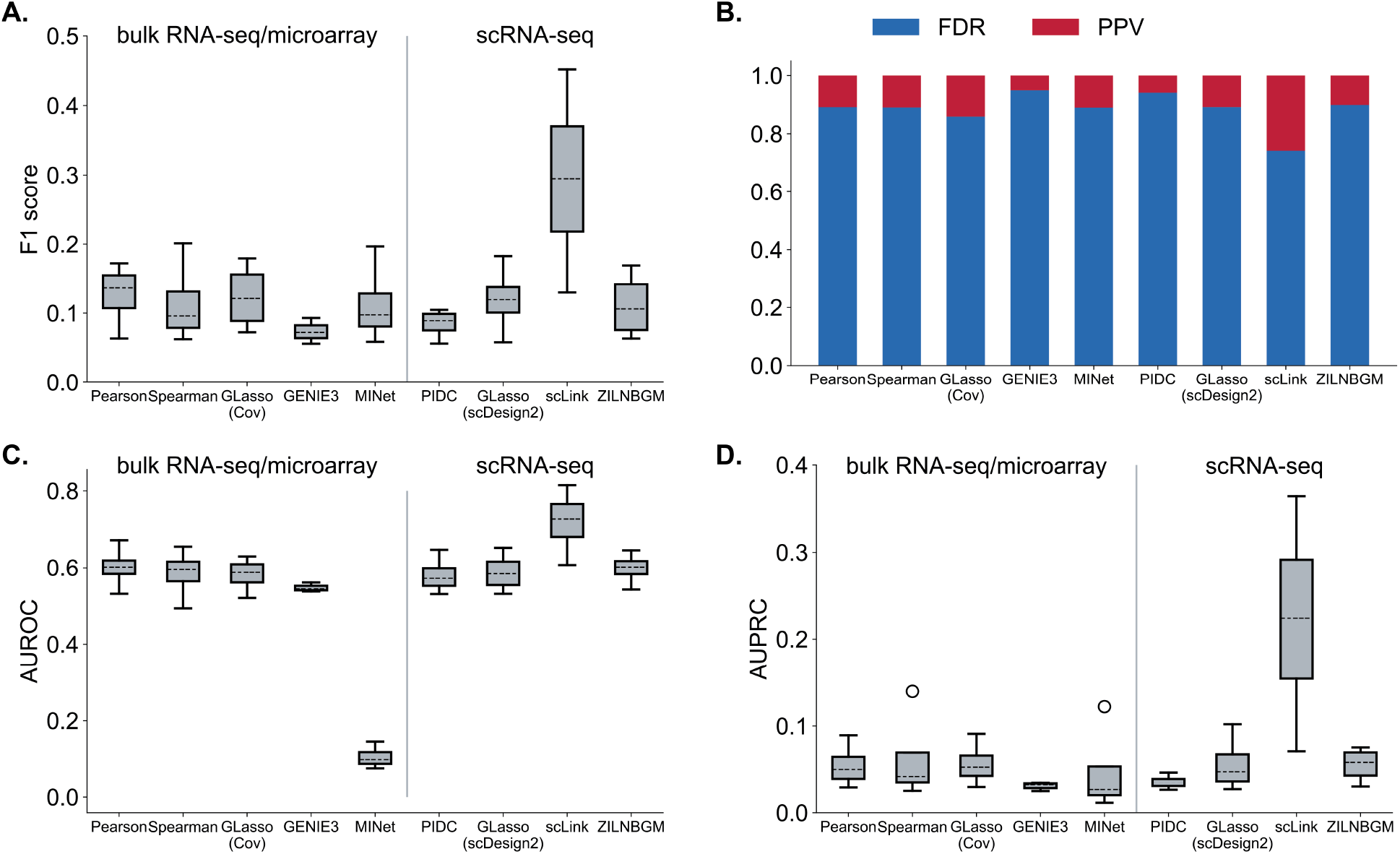
Method performances on four `ZI-Gaussian` simulations with 100 genes, including (**A**) boxplot for F1-score, (**B**) FDR and PPV, (**C**) boxplot for AUROC, and (**D**) boxplot for AUPRC.

**Figure A10:**
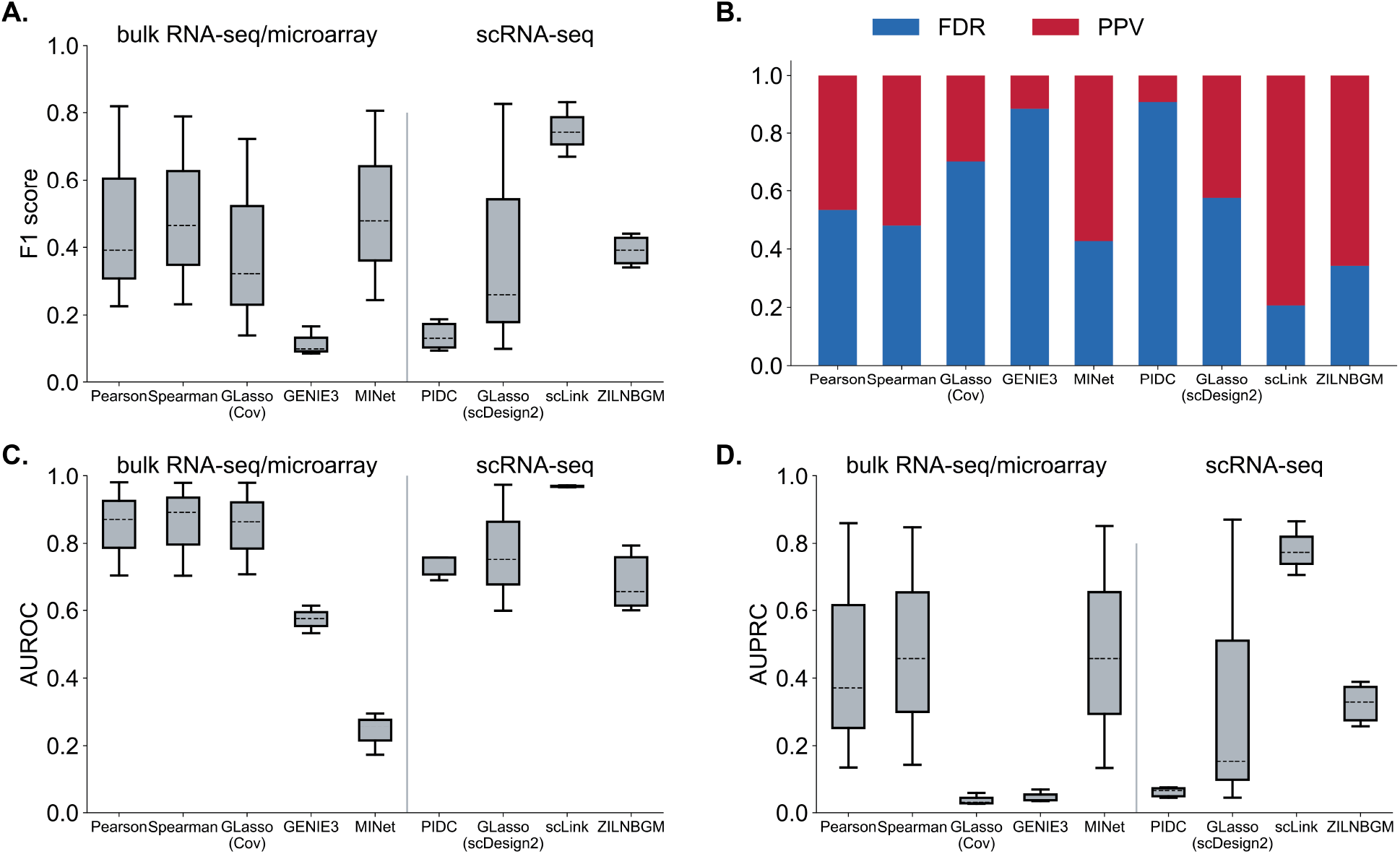
Method performances on three `ZI-Poisson` simulations with 100 genes, including **A**) boxplot for F1-score, **(B)** FDR and PPV, (**C**) boxplot for AUROC, and (**D**) boxplot for AUPRC.

**Figure A11:**
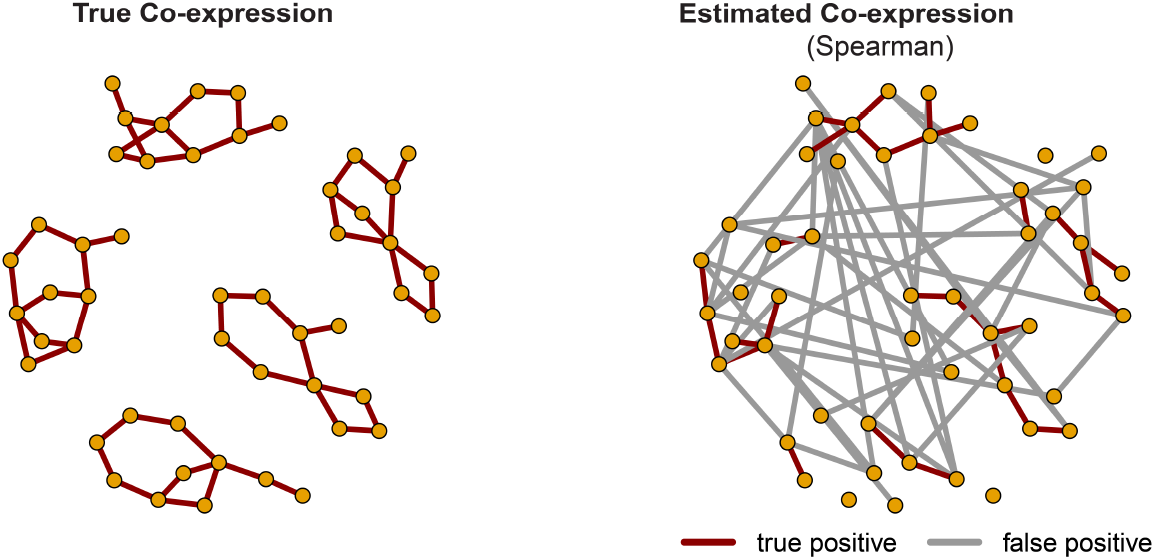
The best `Spearman` estimation on `NORTA` simulations with 100 genes **(right)** and the corresponding true network **(left)**. Here, F1-score=0.43 and FDR=0.6, containing 27 true positives and 40 false positives.

**Figure A12:**
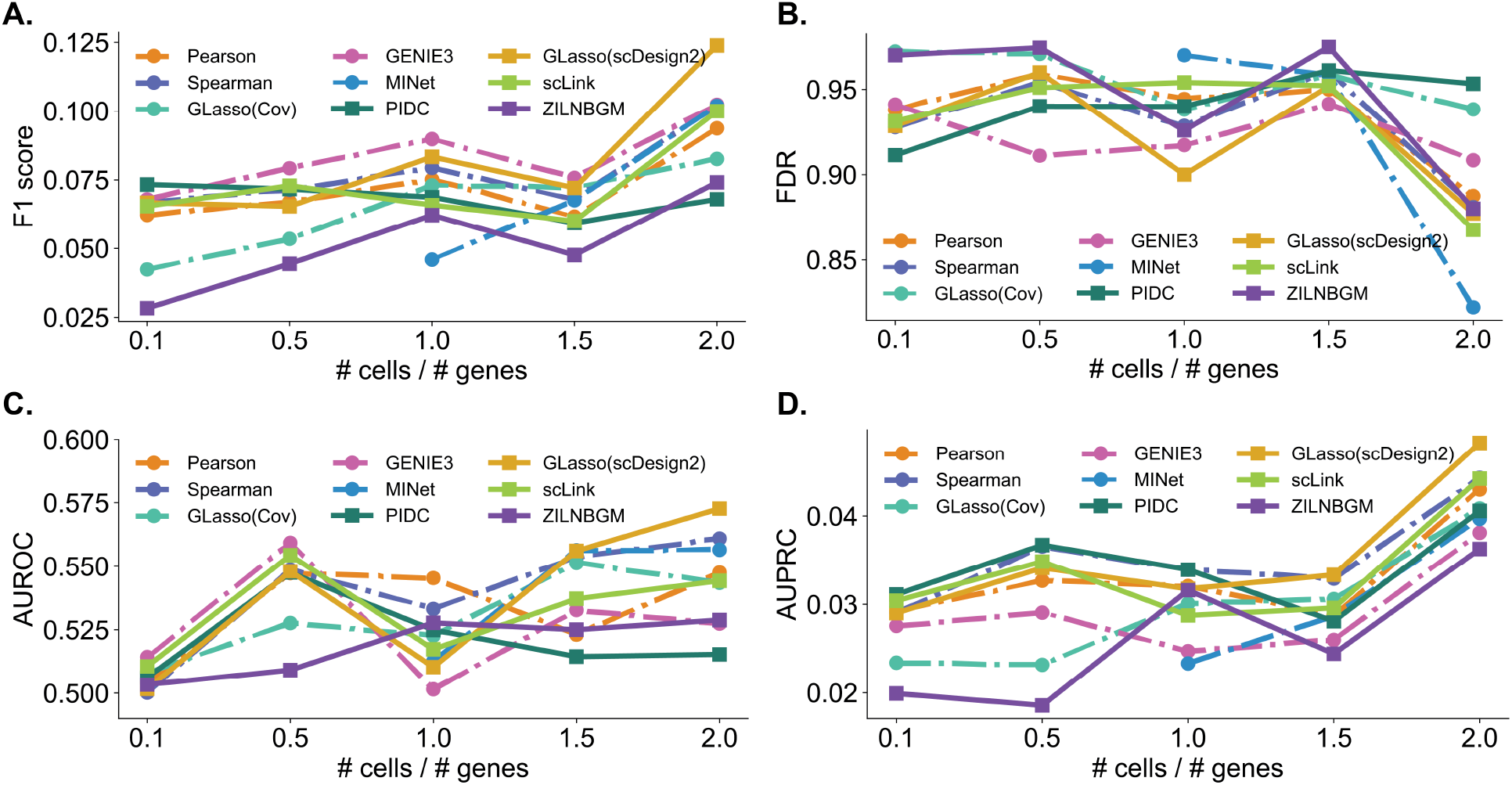
Method performances on `NORTA` simulations with different number of cells and 100 genes. We simulate data with 10, 50, 100, 150, and 200 cells. Evaluations include (**A**) boxplot for F1-score, (**B**) FDR and PPV, (**C**) boxplot for AUROC, and (**D**) boxplot for AUPRC.

**Figure A13:**
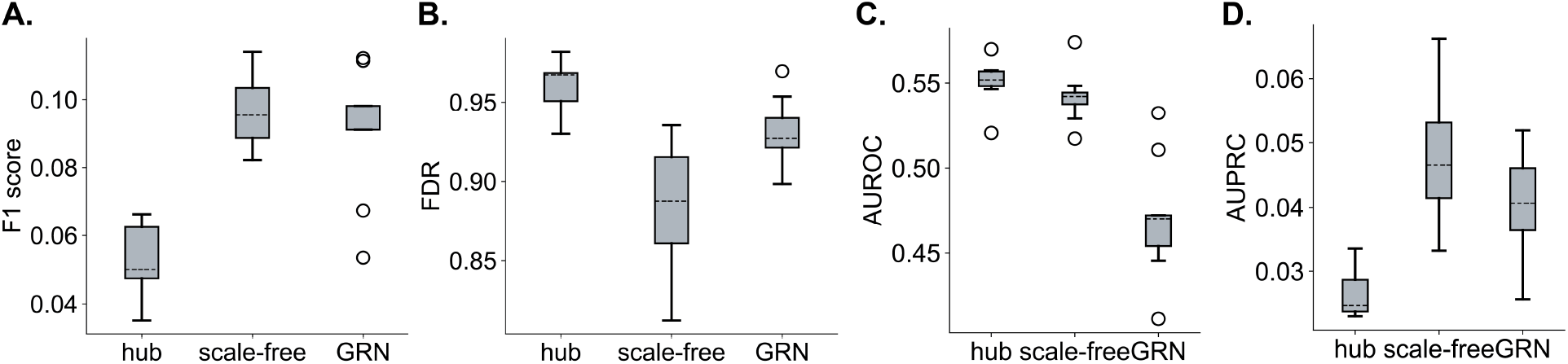
Method performances on `NORTA` simulations with different graph structures and 100 genes. We simulate data with hub-based, scale-free, and regulatory networks. Evaluations include (**A**) boxplot for F1-score, (**B**) FDR and PPV, (**C**) boxplot for AUROC, and (**D**) boxplot for AUPRC.

**Figure A14:**
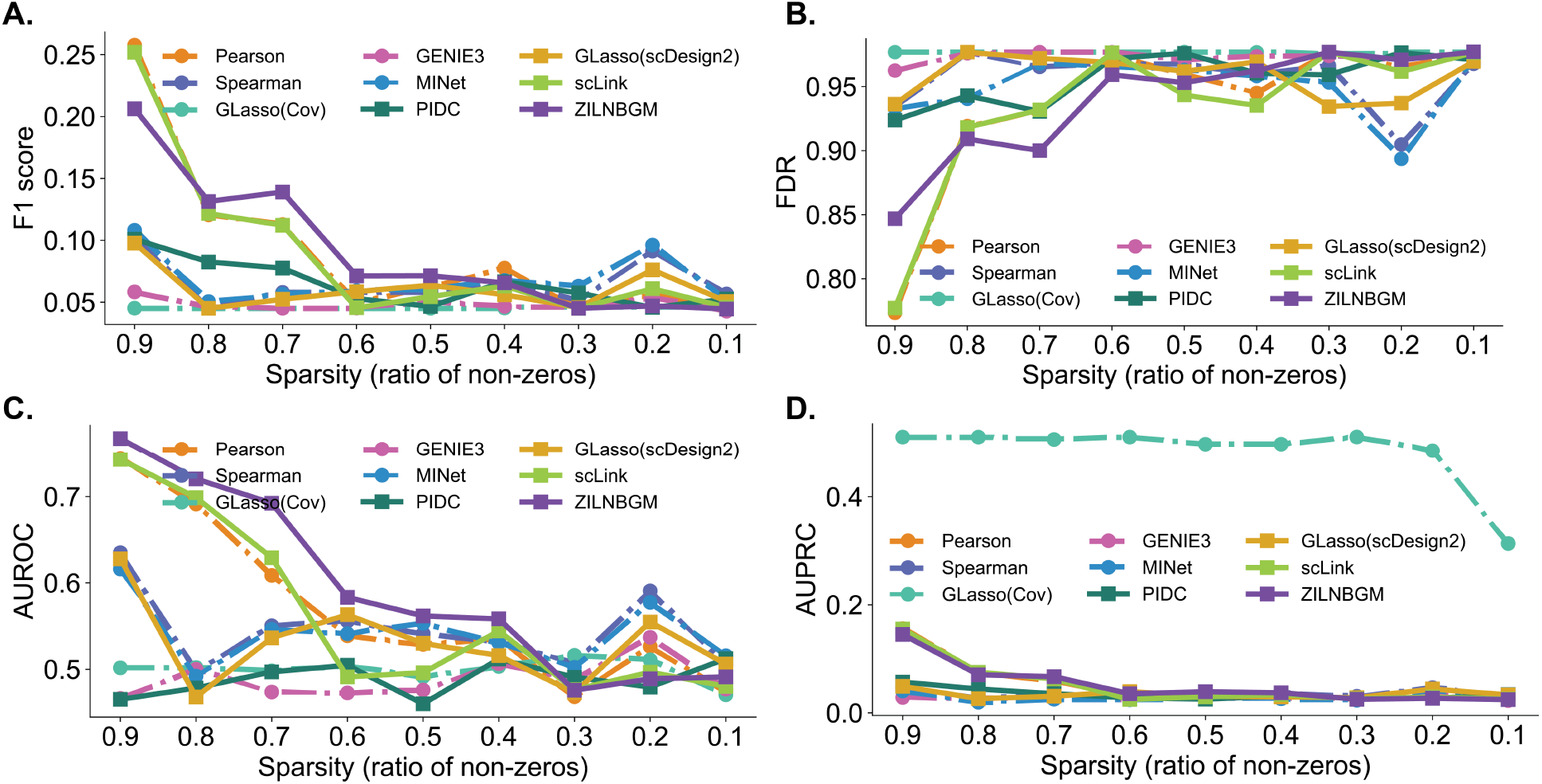
Method performances on `NORTA` simulations with different sparsity levels and 100 genes. We simulate data with 0.1, 0.2, …, 0.9 ratio of non-zero values. Evaluations include (**A**) boxplot for F1-score, (**B**) FDR and PPV, boxplot for AUROC, and (**D**) boxplot for AUPRC.

**Figure A15:**
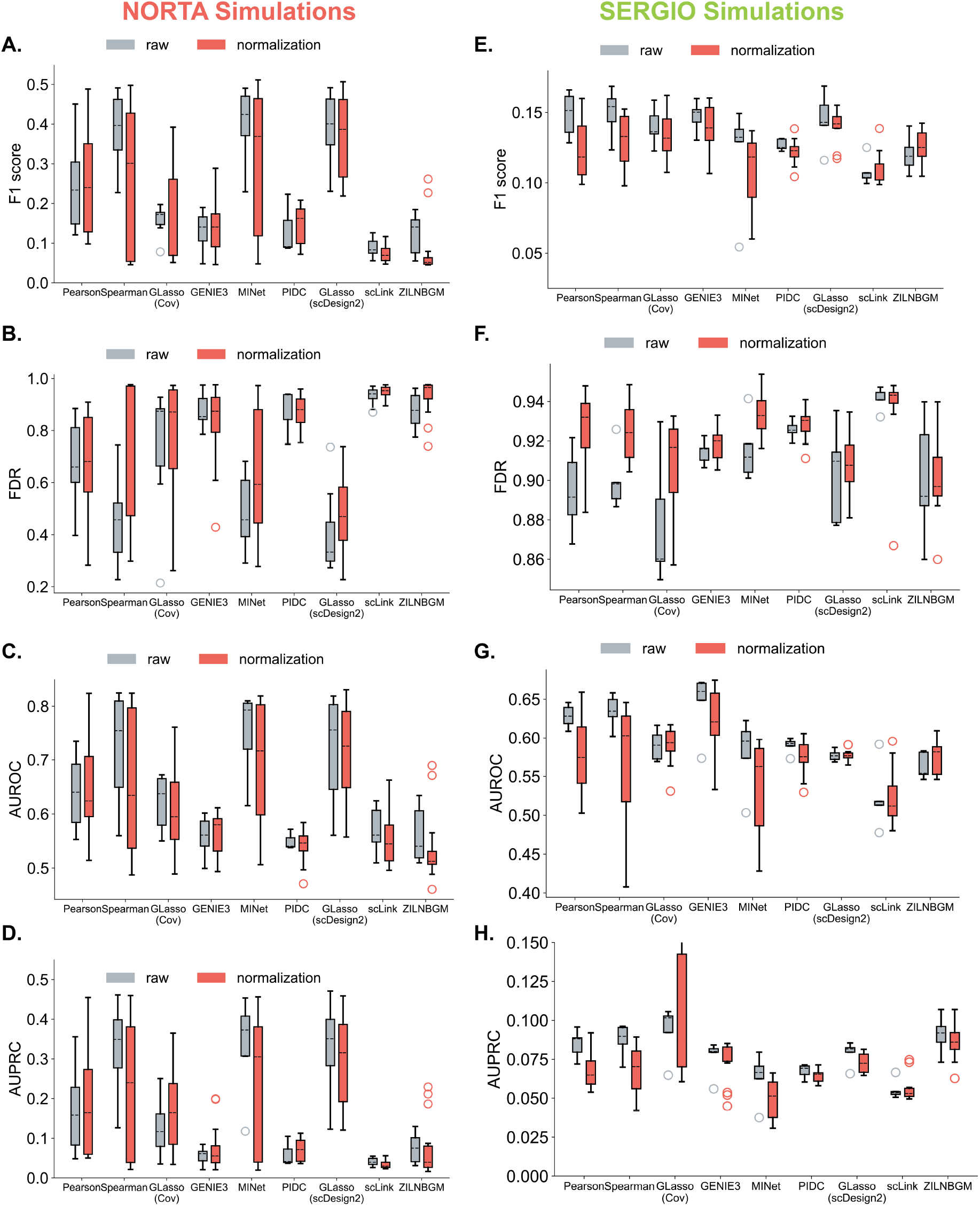
Comparisons of method performances between raw and noramalized `NORTA` (*left column*) and `SERGIO` (*right column*) simulations. Evaluations include (**A**,**E**) boxplot for F1-score, (**B**,**F**) FDR and PPV, (**C**,**G**) boxplot for AUROC, and (**D**,**H**) boxplot for AUPRC.

**Figure A16:**
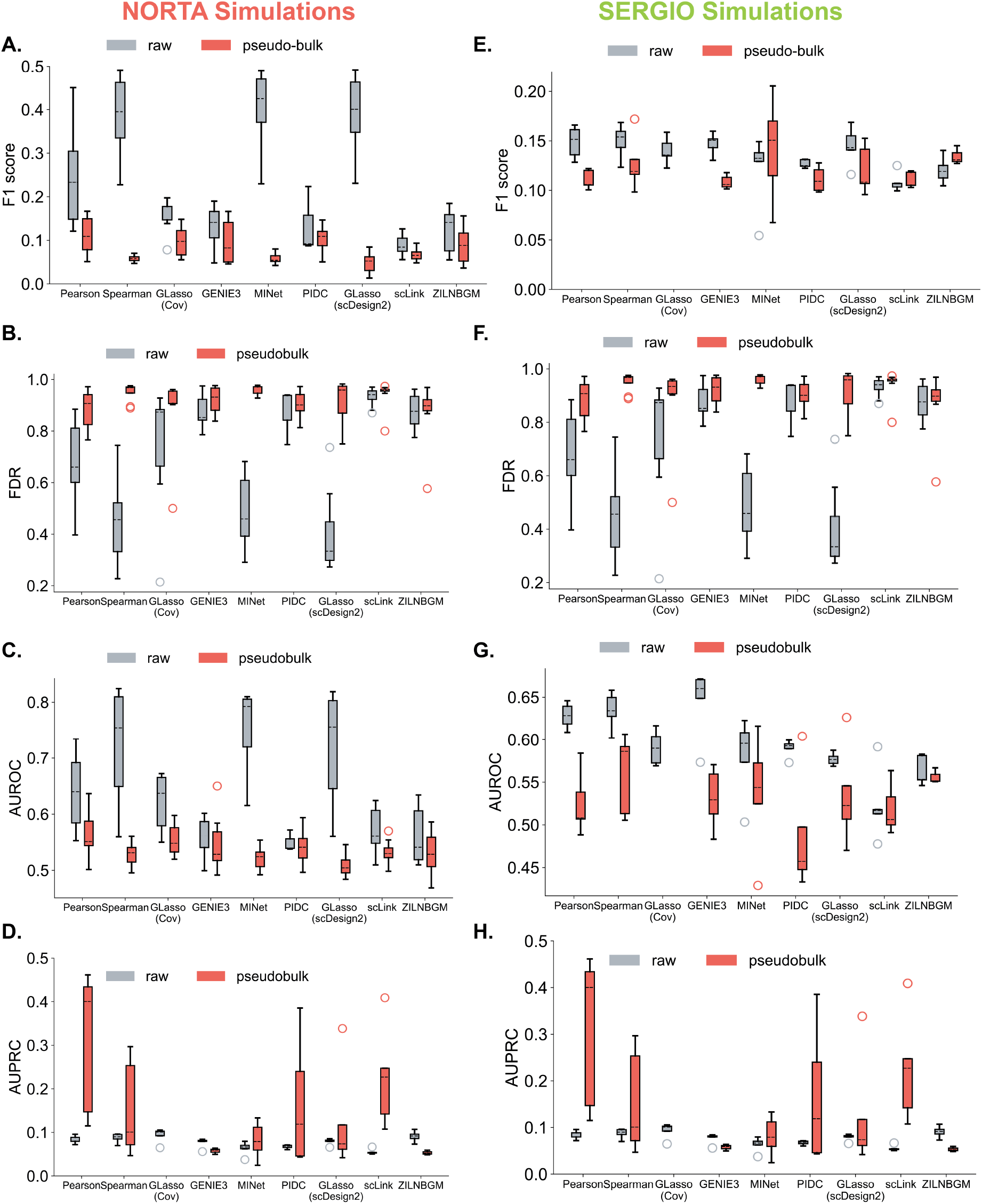
Comparisons of method performances between raw data and pseudo-bulks of `NORTA` (*left column*) and `SERGIO` (*right column*) simulations. Evaluations include (**A**,**E**) boxplot for F1-score, (**B**,**F**) FDR and PPV, (**C**,**G**) boxplot for AUROC, and (**D**,**H**) boxplot for AUPRC.

**Figure A17:**
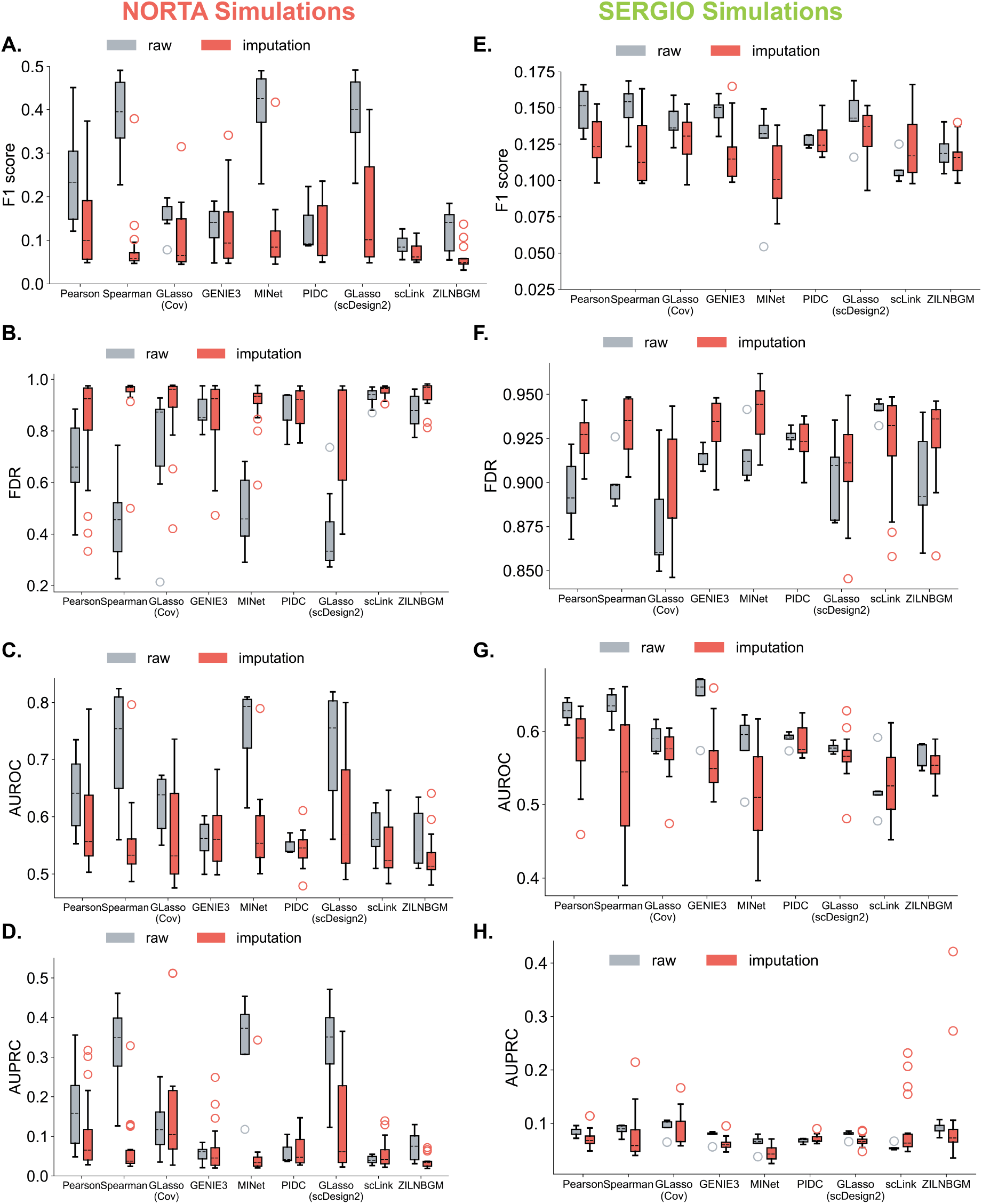
Comparisons of method performances between raw and imputed `NORTA` (*left column*) and `SERGIO` (*right column*) simulations. Evaluations include ((**A**,**E**) boxplot for F1-score, (**B**,**F**) FDR and PPV, (**C**,**G**) boxplot for AUROC, and (**D**,**H**) boxplot for AUPRC.

**Figure A18:**
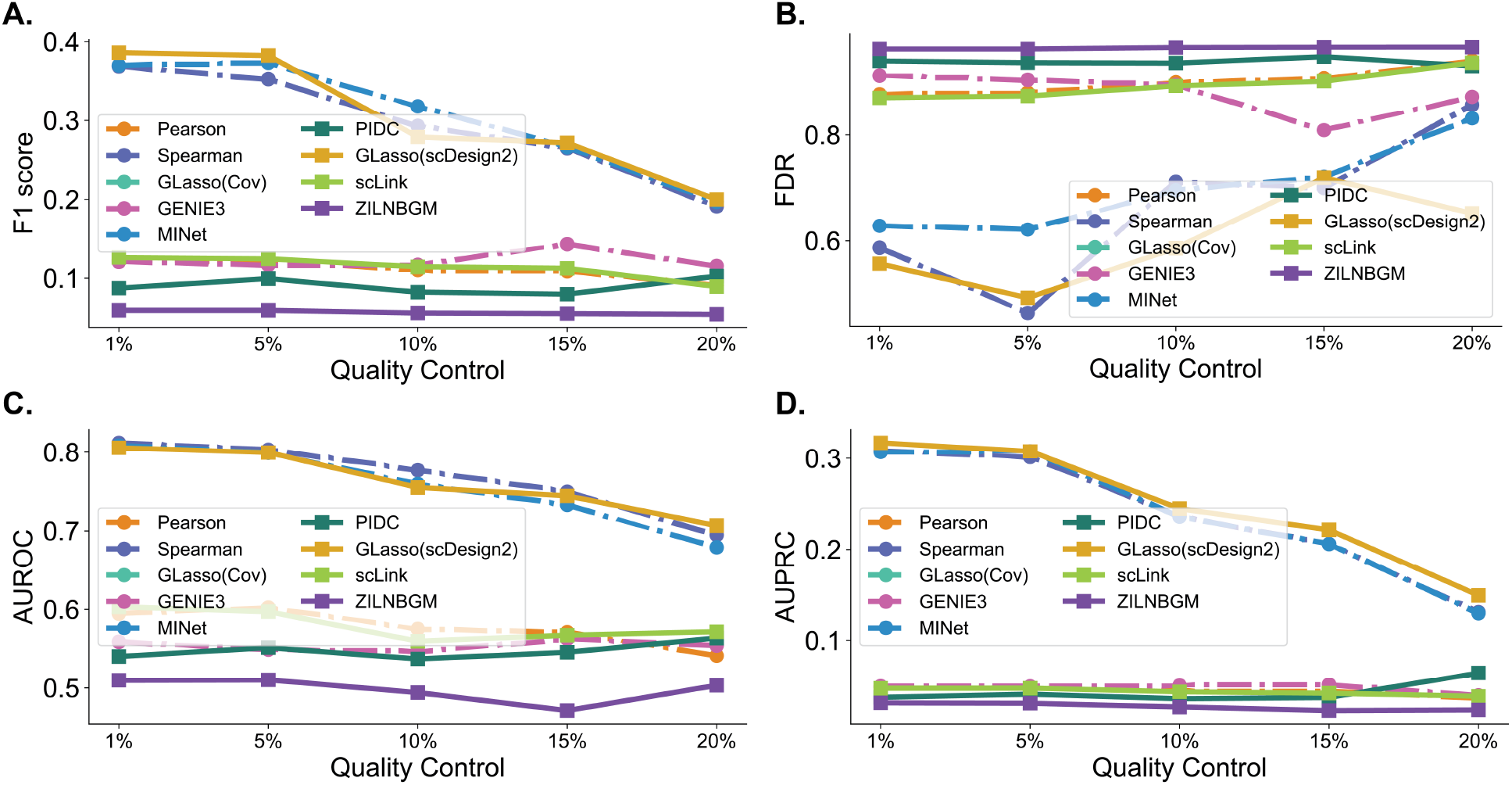
Method performances of `NORTA` simulations after removing cells in the quality control. We first found the maximal library size of all celss and denoted it as *M*. Then for each `NORTA` simulation dataset, we remove cells with its library size smaller than the 1%, 5%, 10%, 15%, and 20% of *M*. Evaluations include **A**) boxplot for F1-score, (**B**) FDR and PPV, (**C**) boxplot for AUROC, and (**D**) boxplot for AUPRC.

**Figure A19:**
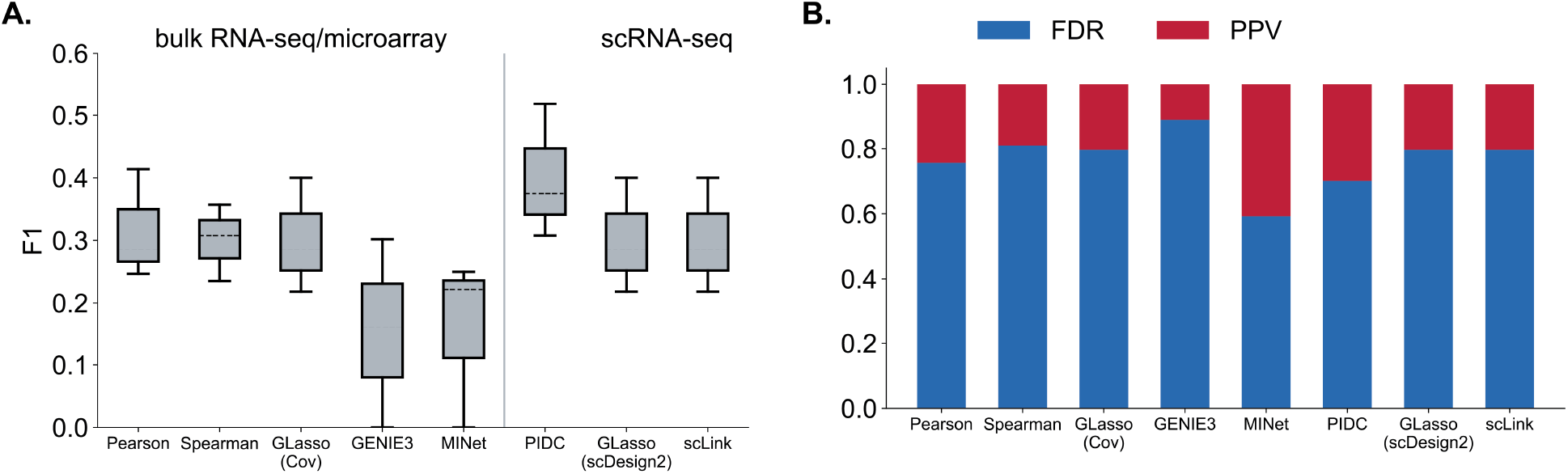
We test methods on Tabula Muris data. We use the top 500 HVGs of three cell types: T cells, pancreatic beta cells, and skeletal muscle satellite stem cells. Because `ZILNBGM` spends several days running a single trial on these data, we consider it uncomparable with other methods and omit it in this comparison.

## Footnotes

2 ℤ* denotes the non-negative integers set.

3 https://singlecell.broadinstitute.org/single_cell. Mouse cortex data study number is SCP425; human PBMC data study number is SCP424.

4 We use the `rnorta` function in `SimCorMultRes` (Touloumis, 2016) R packages in experiment implementations.

5 In the network generation (Sec. 2.1.2), the **G**\* can have negative co-expression. But here, to simplify the evaluation, we only focus on the existence or absence of co-expression.

6 Here, we use 1000 HVGs instead of 100 HVGs because the 100 HVGs contains only a few TFs such that the reference network will be too small.

